# Mate discrimination among subspecies through a conserved olfactory pathway

**DOI:** 10.1101/854364

**Authors:** Mohammed A. Khallaf, Thomas O. Auer, Veit Grabe, Ana Depetris-Chauvin, Byrappa Ammagarahalli, Dan-Dan Zhang, Sofía Lavista-Llanos, Filip Kaftan, Jerrit Weißflog, Luciano M. Matzkin, Stephanie M. Rollmann, Christer Löfstedt, Aleš Svatoš, Hany K.M. Dweck, Silke Sachse, Richard Benton, Bill S. Hansson, Markus Knaden

## Abstract

Signaling mechanisms underlying the sexual isolation of species are poorly understood. Using four subspecies of *Drosophila mojavensis* as a model, we identify two behaviorally active male-specific pheromones. One functions as a conserved male anti-aphrodisiac in all subspecies and acts via gustation. The second induces female receptivity via olfaction exclusively in the two subspecies that produce it. Genetic analysis of the cognate receptor for the olfactory pheromone indicates an important role for this sensory pathway in promoting sexual isolation of subspecies, in collaboration with auditory signals. Surprisingly, the peripheral sensory pathway detecting this pheromone is conserved molecularly, physiologically and anatomically across subspecies. These observations imply that subspecies-specific behaviors arise from differential interpretation of the same peripheral cue, reminiscent of sexually conserved detection but dimorphic interpretation of male pheromones in *D. melanogaster*. Our results reveal that, during incipient speciation, pheromone production, detection and interpretation do not necessarily evolve in a coordinate manner.

## Introduction

One of the central questions in evolutionary biology is how populations within a species split to form new species (Dobzhansky, 1937). Divergence in sexual signaling – male traits and female preferences (Lande, 1981; Ritchie, 2007) – has been proposed as one of the significant forces that results in reproductive isolation (Mayr, 1942). This divergence is more pronounced in allopatric populations as a by-product of ecological adaptation to different environments (Endler, 1992; Nosil, 2012; Seehausen et al., 2008). However, despite numerous documented interspecific sexual traits and preferences in animals, the genetic and neural correlates of their evolution remain elusive (Arguello and Benton, 2017; Smadja and Butlin, 2009).

Sexual traits of drosophilid flies differ quantitatively and qualitatively between species (i.e., act as pre-mating isolation barriers) (Spieth, 1952, 1974), and therefore represent attractive models to determine the genetic basis of phenotypic evolution. Flies identify consubspecific mating partners through integration of different sensory modalities such as vision, audition, olfaction, and gustation (Markow and O’Grady, 2005), which promote courtship of appropriate mates and inhibit courtship of inappropriate partners. Insights have been gained regarding the evolution of different sensory modalities that relate to the interspecific variations in male traits or female perception among drosophilids (e.g., vision (Manoli et al., 2005), audition (Ding et al., 2016), and taste (Ahmed et al., 2019; Seeholzer et al., 2018)). However, even though many studies have reported the diversity of volatile pheromones in drosophilid males (Symonds and Wertheim, 2005), the evolution of the corresponding neural changes remains uninvestigated.

In *D. melanogaster*, sex pheromones are detected by chemosensory receptors (e.g., odorant receptors (ORs)) (Auer and Benton, 2016) expressed in sensory neurons housed in hair-like structures called sensilla. Volatile pheromone-responsive ORs represent an ideal set of candidate genes to address the evolutionary basis of various sexual traits in drosophilid flies due to several reasons. First, pheromone ORs are expected to be the fastest evolving chemosensory receptors, with new receptors emerging either by sequence variation or gene loss/duplication (Guo and Kim, 2007), to match the dramatic diversity of pheromones among closely related species (Symonds and Wertheim, 2005). Second, the neural processing of some drosophilid pheromones (e.g., (*Z*)-11-octadecenyl acetate (*cis* vaccenyl acetate, *c*VA) in *D. melanogaster*) in the brain is well understood (Auer and Benton, 2016). Third, pheromone ORs are narrowly tuned to fly odors (van Naters and Carlson, 2007) and govern instant, robust and distinct behaviors via labeled-line circuitry (Dweck et al., 2015; Ejima et al., 2007; Kurtovic et al., 2007; Liu et al., 2011). Fourth, olfactory sensory neurons (OSNs) that express pheromone ORs target sexually-dimorphic pheromone-processing units (i.e., antennal lobe glomeruli) (Kondoh et al., 2003; Kurtovic et al., 2007). Finally, out of 52 OSN classes in *D. melanogaster* (Benton et al., 2009; Fishilevich and Vosshall, 2005; Grabe et al., 2015), only four pheromone OR-expressing OSNs (Or67d, Or47b, Or65a/b/c and Or88a) are localized in a particular sensillum type (trichoid) (Couto et al., 2005). This small number compared to other non-pheromone-detecting OSNs (Couto et al., 2005) makes a comprehensive evolutionary study feasible.

One remarkable drosophilid is *D. mojavensis*, which represents a model of incipient speciation and host adaptation (Matzkin, 2014). This species has four geographically-isolated and ecologically-distinct populations (Etges, 2019) (taxonomically classified as subspecies (Pfeiler et al., 2009)) that diverged −0.25 million years ago (Etges, 2019; Matzkin, 2014) (Figure 1A). The northern subspecies *D, moj. wrigleyi* and *D. moj. mojavensis* use prickly pear cacti in Santa Catalina Island and red barrel cactus in the Mojave Desert, respectively, as host fruit. The southern subspecies *D. moj. sonorensis* and *D. moj. baja* breed and feed on the organ pipe cactus in the mainland Sonoran Desert and the agria cactus in the Baja California, respectively (Heed, 1978; Ruiz et al., 1990). Phylogenetic analyses revealed that the two northern subspecies and the two southern subspecies clustered with each other (Figure 1B) (Allan and Matzkin, 2019). These closely-related subspecies exhibit many differences in their morphology (Pfeiler et al., 2009), neurophysiological responses (Crowley-Gall et al., 2016; Date et al., 2013; Nemeth et al., 2018), genomic and transcriptomic characteristics (Allan and Matzkin, 2019; Matzkin, 2014; Matzkin and Markow, 2013), and behavioral traits (Newby and Etges, 1998). Experimental reciprocal crosses between these allopatric subspecies resulted in viable fertile offspring, indicating presence of prezygotic isolation mechanisms (Knowles and Markow, 2001; Krebs and Markow, 1989; Markow, 1991; Zouros and Dentremont, 1980). Previous evidence suggested that cuticular hydrocarbons contribute to this isolation barrier (Etges and Ahrens, 2001), but no specific chemicals have been isolated. Here we identify these pheromones and elucidate the evolution of the underlying sensory mechanisms across subspecies to reveal the neurogenetic basis of incipient speciation.

**Figure 1.**
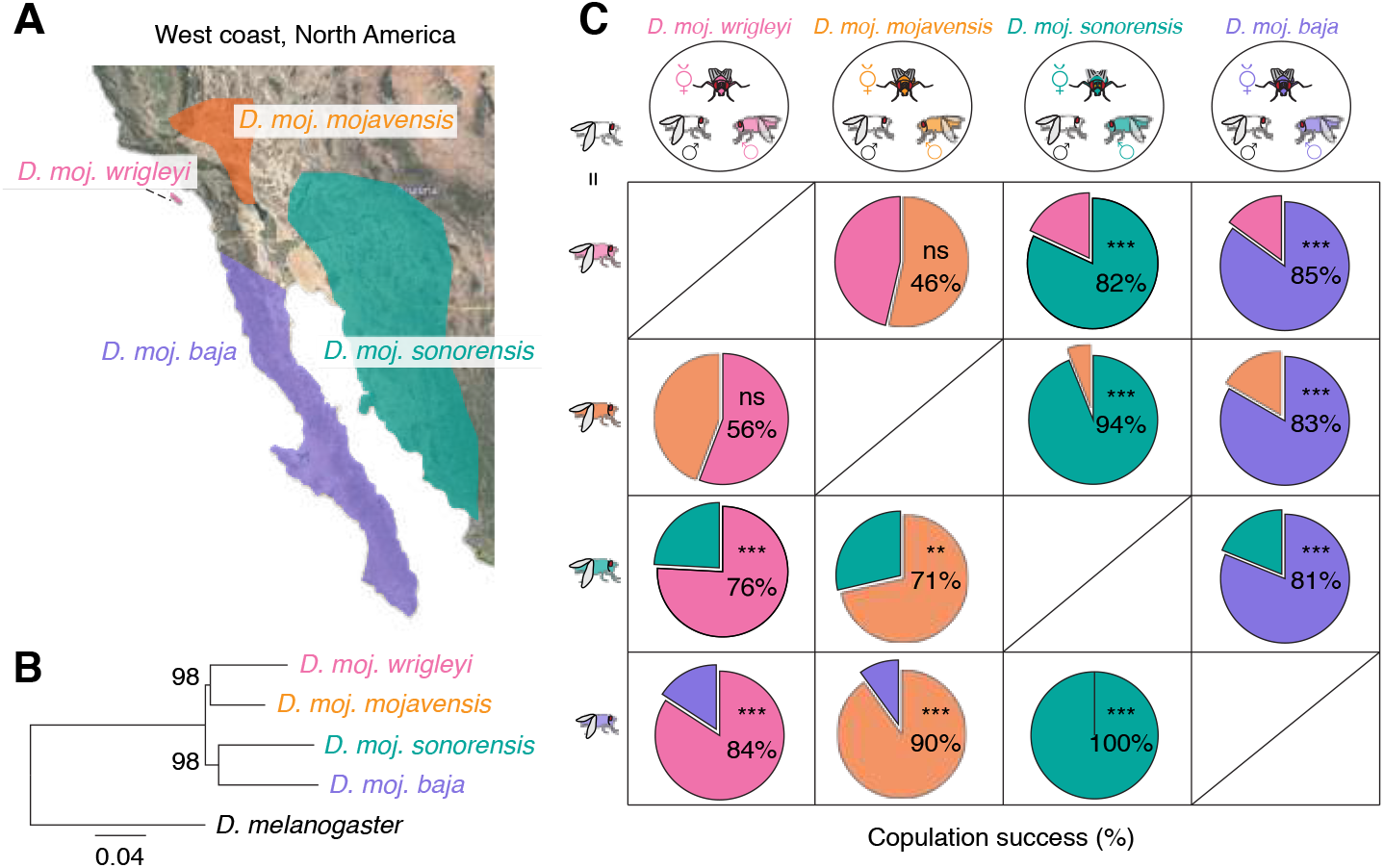
Sexual isolation among *D. mojavensis* subspecies. **A**, Geographic distribution of *D. mojavensis* subspecies on the west coast of North America. Pink, *D. moj. wrigleyi;* orange, *D. moj. mojavensis;* turquoise, *D. moj. sonorensis;* violet, *D. moj. baja;* adapted from (Pfeiler et al., 2009). **B**, Phylogenetic relationship of *D. mojavensis* subspecies based on concatenated sequences of 9087 genes available from (Allan and Matzkin, 2019) (See STAR Methods for details). Scale bar for branch length represents the number of substitutions per site. Bootstrap values are indicated by the numbers at the nodes. **C**, Top row: Competition mating arenas where a female of each *D. mojavensis* subspecies had the choice to mate with a consubspecific male or a male of one of the other three subspecies (color coded as in Figure 1A). Below: Pie-charts represent the percentages of copulation success of the rival males. ns *P* > 0.05; ** *P* < 0.01; *** *P* < 0.001, chi-square test, n=16-25 assays. See Figure S1A for details regarding the differences and similarities of sexual behaviors among the four subspecies.

## Results

### Sexual isolation among incipient subspecies of *D. mojavensis*

We compared the courtship rituals among the four subspecies of *D. mojavensis* by recording the sexual behaviors of consubspecific couples in a single-pair courtship arena (Movie S1-4). *D. mojavensis* subspecies displayed comparable courtship rituals including a novel trait – dropping behavior – in which males release a fluidic droplet from their anus while licking the female’s genitalia (Figure S1).

We asked whether the females of the different subspecies are able to distinguish their consubspecific males by offering a female of each subspecies the choice to mate with a consubspecific male or a male of one of the other three subspecies (Figure 1C). Competition experiments between the two northern subspecies revealed that females did not distinguish consubspecific and hetersubpecific males (Figure 1C). By contrast, southern females strongly preferred to copulate with their consubspecific males (Figure 1C). Both northern and southern females efficiently discriminated northern and southern males (Figure 1C). These results suggest that while a complete sexual isolation barrier has been established both between the northern and southern subspecies, and between the southern subspecies, the northern subspecies are not yet completely sexually isolated.

### *D. mojavensis* subspecies have distinct, but overlapping, profiles of candidate pheromones

To identify candidate pheromones that could mediate the observed sexual isolation barriers between subspecies, we analyzed the chemical profiles of males, and mated and virgin females of each subspecies. We discovered four previously unknown male-specific acetates (Figure 2A and Figure S2A) as *(R)* and (*S*) enantiomers of (*Z*)-10-heptadecen-2-yl acetate (R&S-HDEA), heptadec-2-yl acetate (HDA) and (*Z,Z*)-19,22-octacosadien-1-yl acetate (OCDA). Males of all four subspecies carried OCDA, while only males of the northern subspecies produced HDEA (both *R* and *S* enantiomers in similar ratio (Figure S2B-C)) and HDA (Figure 2A and Figure S2A). The chemical profiles of the virgin females were similar among the different subspecies, whereas mated females carried male-specific compound(s), indicating that these are transferred from males to females during mating (Figure 2A and Figure S2A). Using matrix-assisted laser desorption/ionization-time of flight (MALDI-TOF) (Kaftan et al., 2014) analyses to directly visualize pheromones on the fly body (see STAR Methods for details), we confirmed the existence of OCDA on the male and mated female surfaces, but not on the virgin female (Figure S2D-E).

**Figure 2.**
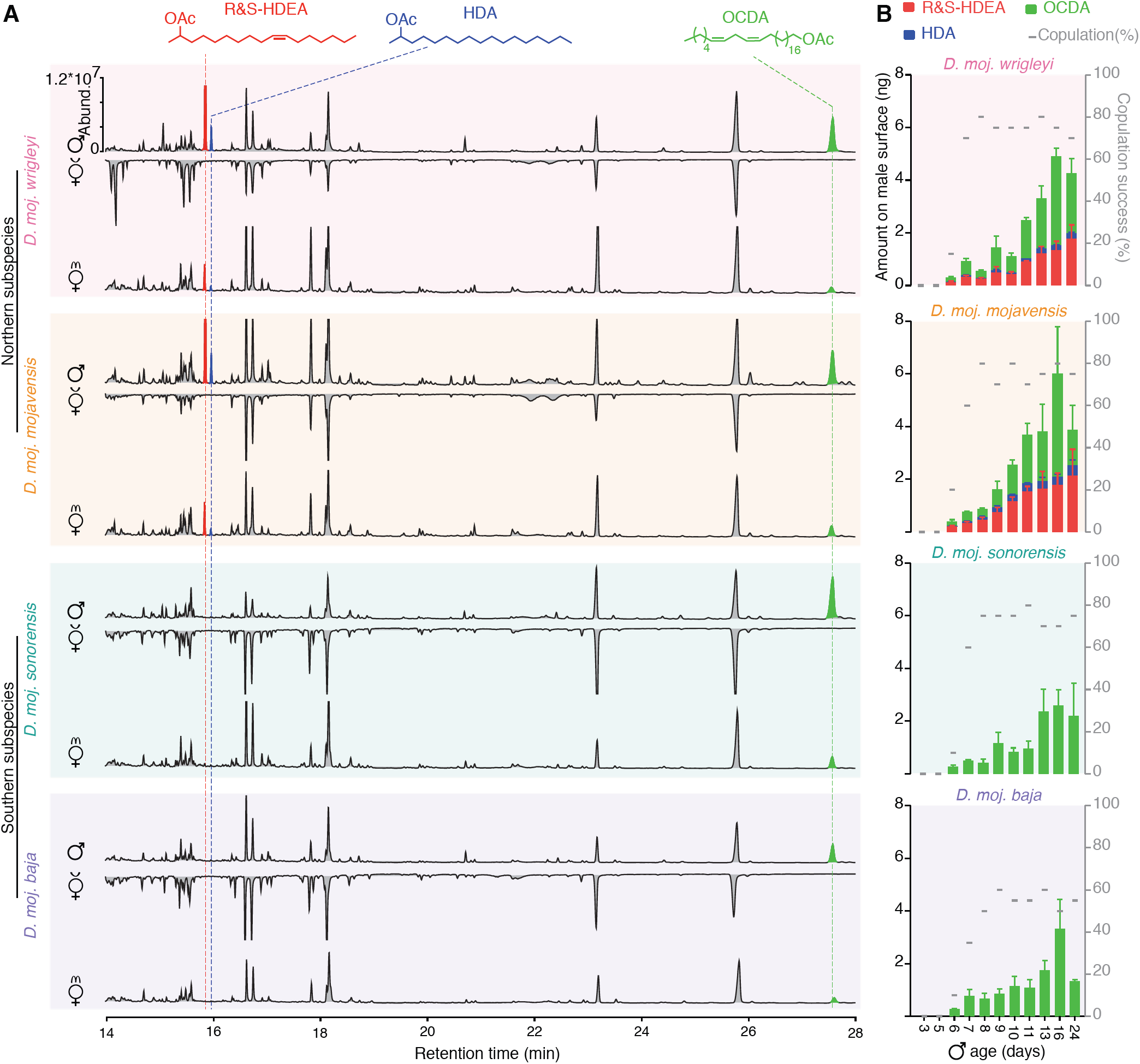
Dissimilar male-specific compounds among *D. mojavensis* subspecies. **A**, Representative gas chromatograms of 10 day old male (virgin, ♂) and female flies (virgin; v♀ and mated; m♀) (n=7) obtained by solvent-free thermal desorption-gas chromatography-mass spectrometry (TD-GC-MS) (Dweck et al., 2015). Colored peaks indicate the male-specific compounds (see STAR Methods for chemical syntheses), which are transferred to females during mating. Red: R and S enantiomers of (*Z*)-10-Heptadecen-2-yl acetate (R&S-HDEA); blue: heptadec-2yl acetate (HDA); light green: (*Z,Z*)-19,22-octacosadien-1yl acetate (OCDA). Different colored backgrounds represent the different subspecies (similar to Figure 1A). In this and other panels *D. moj. wrigleyi* (15081-1352.22), *D. moj. mojavensis* (15081-1352.47), *D. moj. sonorensis* (15081-1351.01), and *D. moj. baja* (15081-1351.04) were used. Two more strains of each subspecies (see Key Resources Table for details) were analyzed showing very similar profiles (data not shown). **B**, Amount of the male-specific compounds and corresponding copulation performance. Colored bars and error bars indicate mean amounts and SEM of the three male-specific acetates (n=5 males per age); grey dashes indicate the percentage of copulation success for males of the same age within a 10-minute time window (n=25 males per age). See Figure S2A for details regarding the body wash extracts analyzed by GC-MS and Figure S2F for the production site of these male-specific compounds.

We next investigated whether production of the male-specific compounds correlates with male sexual maturity and copulation performance. Chemical analysis of male profiles from 0 to 24 post-eclosion individuals indicated that males of the four subspecies exhibit high abundance of the acetate(s) together with an increased copulation success from the 7^th^ day on (Figure 2B). We conclude that the identified male-specific acetates define the maturity status of males.

Taken together, consistent with the phylogenetic relationships (Figure 1B), the two northern subspecies are chemically similar to each other and differ from the two southern subspecies. We identified OCDA in all four *D. mojavensis* subspecies, while R&S-HDEA and HDA are exclusively produced in northern subspecies.

### The ubiquitous pheromone, OCDA, acts as conserved male anti-aphrodisiac via contact chemosensation

We asked if transfer of any male-specific compounds contributes to a general post-copulation mate-guarding strategy in *D. mojavensis* (as described for transferred pheromones in *D. melanogaster*(Yewetal., 2009; Zawistowski and Richmond, 1986)). The identical chemical profiles of the northern subspecies and of the southern subspecies led us to focus our attention on one representative for each group. Consistent with a role of male-transferred compounds during mating, males of *D. moj. wrigleyi* and *D. moj. sonorensis* spent more time courting virgin than mated consubspecific females (Figure 3A). To test which, if any, of the identified compounds contribute to courtship reduction, we perfumed virgin consubspecific females with one of the four male-specific compounds. Only OCDA treatment of females led to a significantly reduced male courtship index (Figure 3B and Figure S3A).

**Figure 3.**
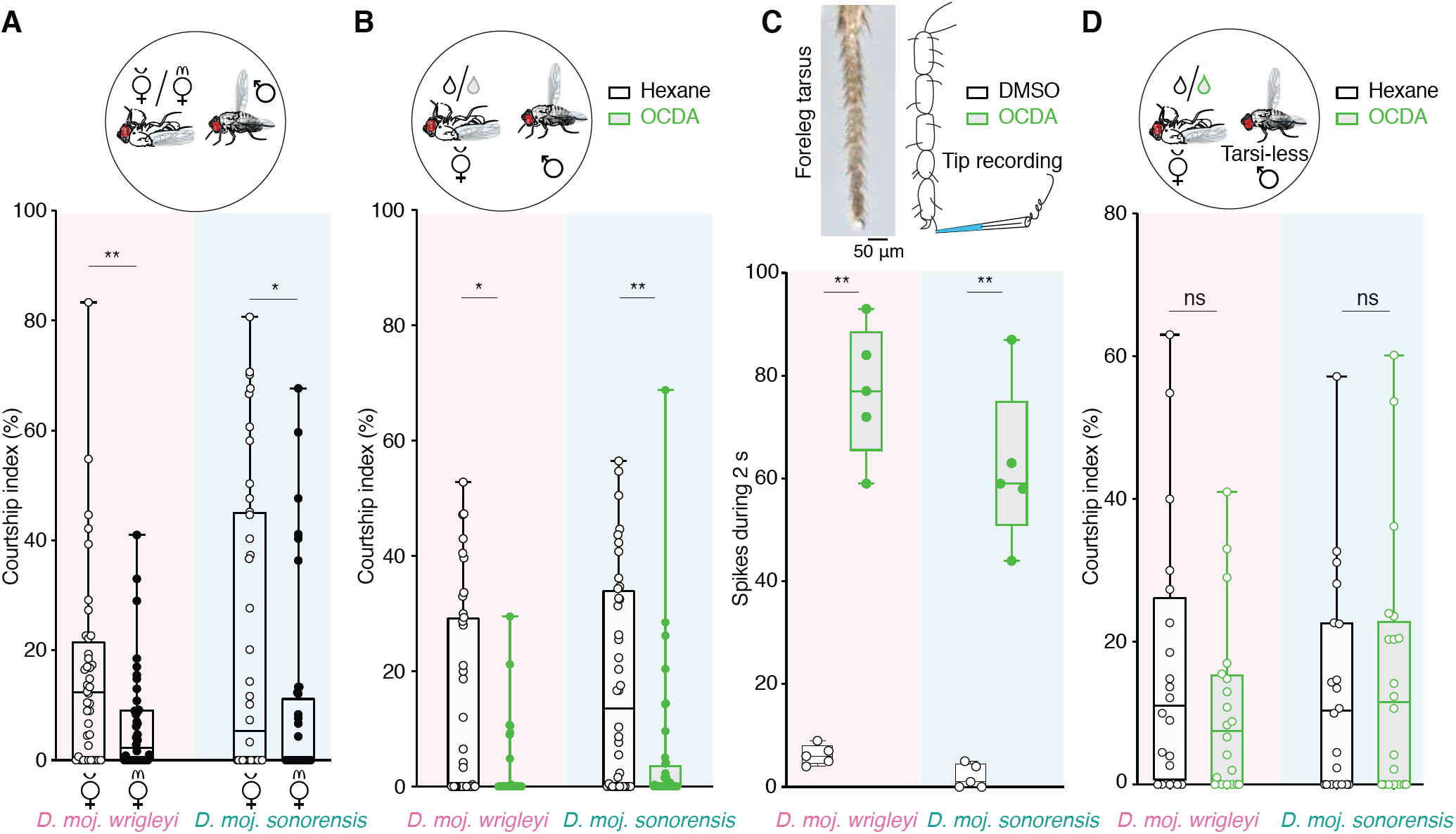
Conserved behavioral responses among *D. mojavensis* subspecies to OCDA. **A**, Top: Schematic of courtship arena where a dead (virgin or mated) female (*D. moj. wrigleyi* and *D. moj. sonorensis)* was presented to a consubspecific male. Below, y-axis represents courtship index [%] (equal the time a male exhibits courtship behaviors (Figure S1A) / total amount of recording time (10 minutes)). In this and other panels, filled circles indicate significant difference between the tested groups; * *P* < 0.05; ** *P* < 0.01, Mann Whitney U test, n=40. Males and females used in this and other panels are 10 days old. **B**, Top: Schematic of a male courting a dead virgin female of the same subspecies perfumed with hexane as a control (black) or OCDA diluted in hexane (light green). Kruskal-Wallis test with Dunn’s post-hoc correction. Ns *P* > 0.05; * *P* < 0.05; ** *P* < 0.01, n=40 assays. See Figure S3A for the role of the other male-specific acetates in male courtship suppression in both subspecies. **C**, Left top: micrograph of tarsal segments of the foreleg showing different sensory hairs. Scale bar = 50 μm. Right top: schematics of tip recordings from the foreleg tarsal sensillum (class 5b). Bottom: tip recording measurements from foreleg-tarsi of *D. moj. wrigleyi* and *D. moj. sonorensis* males using DMSO or OCDA. Kruskal-Wallis test with Dunn’s post-hoc correction. Ns *P* > 0.05; ** *P* < 0.01, n=5. See Figure S3C for tip recording traces and Figure S3D for electrophysiological responses to the other male-specific acetates. **D**, Courtship indices of tarsi-less males tested with a dead virgin female of the same subspecies perfumed with hexane (black) or OCDA (light green). Ns > 0.05, Mann Whitney U test, n=20 assays. See Figures S3E-F’’ for behavioral assays testing the roles of these male-specific acetates in regulating social behaviors (i.e, aggregation) of *D. moj. wrigleyi* and *D. moj. sonorensis*.

To investigate how males detect OCDA, we first tested the volatility of male-specific acetates (including OCDA) by collecting headspace samples of *D. moj. wrigleyi* males (Figure S3B). This revealed that only R&S-HDEA and HDA are volatile (Figure S3B) indicating that the non-volatile OCDA is likely to be detected by gustation. During courtship, neural inputs from foreleg gustatory sensilla are used to evaluate the potential mating partner (Spieth, 1974). We therefore investigated whether male foreleg tarsi detect OCDA (Figure 3C). Indeed, OCDA, but not other acetates, elicited a response in a subset of tarsal sensilla (class 5b) of both *D. mojavensis* subspecies (Figure 3C and Figure S3C-D). Consistent with these sensilla mediating detection of OCDA, males lacking their tarsi spend equal time courting the hexane-perfumed and OCDA-perfumed females (Figure 3D). We conclude that OCDA has a conserved function among the *D. mojavensis* subspecies, is detected by tarsal gustatory sensilla and acts as an anti-aphrodisiac signal that suppresses male courtship.

### R-HDEA promotes subspecies-specific female sexual receptivity through olfaction

To examine the functions of the other novel male-specific acetates in female sexual behaviors, we scored the copulation success and latency for *D. moj. wrigleyi* and *D. moj. sonorensis* females courted by consubspecific males. Males were perfumed with one of the four male-specific acetates or hexane. Copulation success did not differ between the acetate- and hexane-perfumed males in both subspecies (Figure 4A and Figure S4A). However, females of *D. moj. wrigleyi* were significantly quicker to accept the R-HDEA-perfumed males (Figure 4B and Figure S4B). We extended this analysis by performing competition assays, in which a virgin female of each subspecies was allowed to choose between two consubspecific males perfumed with hexane or with one of the four acetates. Only R-HDEA-perfumed males exhibited copulation advantage over the controls in *D. moj. wrigleyi* and *D. moj. mojavensis*, while *D. moj. sonorensis* and *D. moj. baja* females had comparable preferences between the two males (Figure 4C and Figure S4C-D). These results indicated that R-HDEA increases sexual receptivity in females of northern but not southern subspecies.

**Figure 4.**
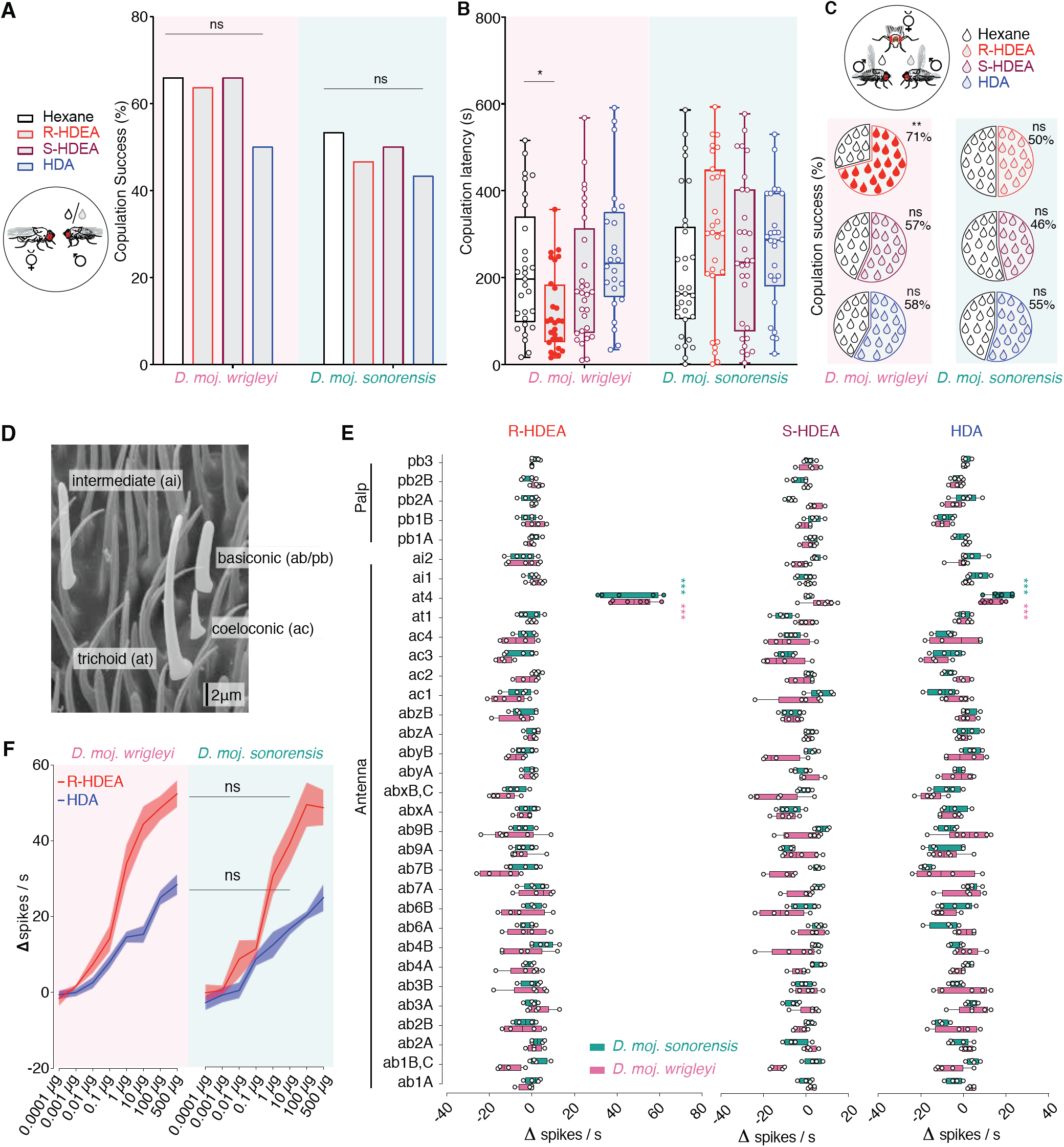
R-HDEA provokes divergent sexual behaviors through activation of homologous sensory neurons. **A**, Left: Schematic of mating arena where a perfumed male courts a virgin consubspecific female. Black droplet: hexane (control); grey droplet: one of the other three male-transferred compounds. Right: Copulation success of *D. moj. wrigleyi* and *D. moj. sonorensis* males perfumed with hexane or male-specific acetates. Fisher exact test. Ns *P* > 0.05, n=40 assays. See Figure S4A for OCDA impact on copulation success. **B**, Copulation latencies of the same males as in Figure 4A. Color-filled circles indicate significant differences from the solvent. Kruskal-Wallis test with Dunn’s post-hoc correction. Ns *P* > 0.05; * *P* < 0.05, n=40 assays. **C**, Competition between two consubspecific males, perfumed with one of the three compounds (colored droplet) or with the solvent hexane (black droplet), to copulate with a virgin female of the same subspecies. Pie-charts represent copulation success of the rival males. Filled droplets indicate significant difference between the tested groups. Chi-square test, Ns *P* > 0.05; ** *P* < 0.01, n=26-31 assays. See Figure S4C for the competition results of OCDA-perfumed males. See Figure S4D for *D. moj. mojavensis* and *D. moj. baja* competition experiments. **D**, Scanning electron micrograph of antennal surface showing different sensillum types (intermediate, trichoid, coeloconic and basiconic). Scale bar = 2 μm. **E**, Electrophysiological responses towards R-HDEA, S-HDEA and HDA in all types of olfactory sensilla on antenna and maxillary palp of *D. moj. wrigleyi* (pink) and *D. moj. sonorensis* (turquoise). Mann Whitney U test, ns *P* > 0.05; *** *P* < 0.001; n=3-6 neurons. ab, antennal basiconic; ac, antennal coeloconic; at, antennal trichoid; ai, antennal intermediate; pb, palp basiconic. See Figure S4E-G for similarity between male and female responses to R-HDEA, OCDA responses, and representative SSR traces, respectively. **F**, Dose-dependent responses of at4 neurons in *D. moj. wrigleyi* and *D. moj. sonorensis* toward R-HDEA (in red) and HDA (in blue) (Mean ± SEM). Two-way ANOVA followed by Sidek’s multiple comparison test between the two subspecies’ responses to the same stimulus, ns P > 0.05; n=4-6 neurons. See Figure S4I for *D. melanogaster* at1 and at4 responses to *D. moj. wrigleyi-* specific acetates.

As R-HDEA is volatile (Figure S3B) we searched via single sensillum recordings (SSR) for olfactory sensory neurons (OSNs) that detect R-HDEA in *D. moj. wrigleyi*. We identified a single antennal trichoid sensillum class, at4 (homologous to the *D. melanogaster* at4 sensillum REF) that responds strongly to R-HDEA and HDA. (Figure 4D-E and Figure S4E-G)). Unexpectedly, *D. moj. sonorensis* has a sensillum with identical response properties (Figure 4E-F and Figure S4E-H) even though this subspecies neither produces nor responds behaviorally towards R-HDEA (Figures 2A and 4B-C). Taken together, R-HDEA activates homologous olfactory channels in all subspecies of *D. mojavensis* and enhances sexual receptivity in *D. moj. wrigleyi* but not in *D. moj. sonorensis* females.

### Conserved peripheral sensory pathways for R-HDEA detection

To define the genetic basis of R-HDEA detection in *D. mojavensis* subspecies, we focused on receptors expressed in trichoid OSNs, which were previously shown to detect pheromones in *D. melanogaster* (Couto et al., 2005; Dweck et al., 2015; Kurtovic et al., 2007; van Naters and Carlson, 2007). Five orthologous genes are present in the *D. moj. wrigleyi* genome (Guo and Kim, 2007): *Or47bl, Or47b2, Or65a, Or67d* and *Or88a* (Figure S5A for terminology and details). We first visualized their expression in the antenna of *D. moj. wrigleyi* and *D. moj. sonorensis* by RNA *in situ* hybridization and detected expression for all receptors with a slightly increased number of *Or65a* and *Or67d-*expressing cells in *D. moj. wrigleyi* (Figure 5A-B). We then asked which of these ORs is responsible for R-HDEA detection by functional expression in *Xenopus laevis* oocytes (Figure 5C) and could detect significant depolarization upon R-HDEA application for OR47b1 and OR65a (Figure 5D and Figure S5B). To validate these results, we generated transgenic flies expressing *D. moj. wrigleyi* ORs individually *in vivo* in the at1-neuron of *D. melanogaster* lacking its endogenous receptor OR67d (Kurtovic et al., 2007) (decoder neuron) (Figure 5E). SSR analysis of these flies revealed that OR65a is the sole detector for R-HDEA and HDA (Figure 5F and Figure S5C). As predicted by the similar responses of at4 sensilla to R-HDEA and OR65a sequences among both subspecies (Figure 4E-G and Figure S5G), transgenic expression of OR65a of *D. moj. sonorensis* in the decoder neuron revealed comparable responses to R-HDEA (Figure 5G). Together, we conclude that *D. moj. wrigleyi* and *D. moj sonorensis* detect R-HDEA via OR65a.

**Figure 5.**
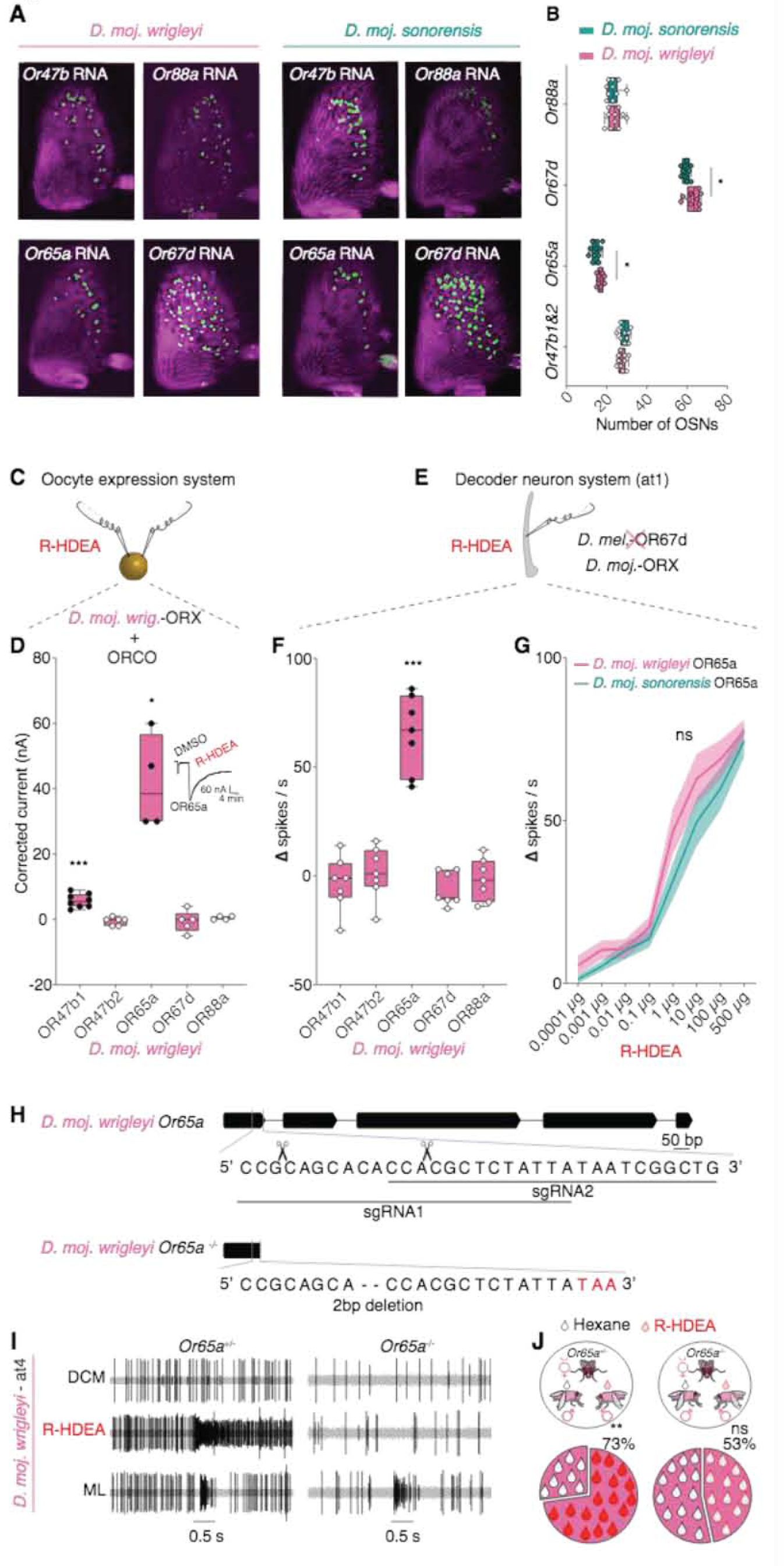
Conserved detection mechanism of R-HDEA among *D. mojavensis* subspecies. **A,** Expression of olfactory receptor genes (*OrX: Or47b, Or88a, Or65a* and *Or67d*) in *D. moj. wrigleyi and D. moj. sonorensis* female antennae. See STAR Methods, Figure S5A and Figure S4J for the receptors’ terminology and relationships. Due to the high degree of sequence identity (99.1%) of the *Or47b-like* loci (File S1), cross-hybridization between probes and mRNAs is likely to happen. **B,** Number of the Or-expressing cells (*OrX: Or47b, Or88a, Or65a* and *Or67d*) in *D. moj. wrigleyi* and *D. moj. sonorensis* females. Color-filled circles indicate significant differences between both species. Mann Whitney U test, ns *P* > 0.05; * *P*< 0.05; n=10 antennae. **C,** Left: Schematic of voltage-clamp recordings from *Xenopus laevis* oocytes ectopically expressing the different odorant receptors (ORX) together with ORCO. Right: Traces of electrophysiological responses of D. moj. wrigleyi-OR65a to DMSO and R-HDEA. **D,** Electrophysiological responses of oocytes expressing different odorant receptor genes to R-HDEA (1mM). Black-filled circles indicate significant differences from the solvent (DMSO) response. Mann Whitney U test, ns *P* > 0.05; * *P* < 0.05; *** *P* < 0.001, n=5-8 recordings. See Figure S5B for details. **E,** Schematic of odorant receptors (OrX) expressed in *D. melanogaster* at1 neurons. **F,** Electrophysiological responses of five odorant receptors expressed in *D. melanogaster* at1 neurons to R-HDEA (10 μg). Black-filled circles indicate significant differences from the solvent (Hexane) response. Mann Whitney U test, ns *P* > 0.05; *** *P* < 0.001, n=7 recordings. See Figure S5B for details. **G,** Electrophysiological responses of *D. moj. wrigleyi* or *D. moj. sonorensis* OR65a towards increasing concentrations of R-HDEA (n=5-8 recordings). Two-way ANOVA followed by Sidek’s multiplecomparison test between the two subspecies’ responses to the same stimulus, ns P > 0.05; n=4-6 neurons. **H**, Schematics of the *D. moj. Or65a* locus illustrating the sgRNA binding sites. Scissors denote the presumed cutting site. The *Or65a* loss-of-function allele carries a 2 bp deletion in exon 1, resulting in a premature stop codon (highlighted in red). **I**, Electrophysiological responses of *Or65a* heterozygous (left) and homozygous (right) animals to hexane, R-HDEA and methyl laurate (ML). The latter shows an intact neuronal excitation of the Or65a-neighbouring neuron. See Figure S5D regarding quantification of SSR responses. **J**, Competition between two males of *D. moj. wrigleyi*, perfumed with R-HDEA (red droplet) or hexane (black droplet), to copulate with a *D. moj. wrigleyi* virgin female. Left: Heterozygous animal at the *Or65a* locus, n = 37; right: homozygous mutant at the *Or65a* locus, n = 32. Colored filled droplets indicate a significant difference between the rival males. Chi-square test, ns *P* > 0.05; ** *P* < 0.01. All males and females used in this and other panels were 10-day-old virgin flies. See Figure 6SD-E for details regarding the *D. moj. wrigleyi white* mutant.

To determine whether *Or65a* is required for the enhanced female receptivity in *D. moj. wrigleyi*, we generated a loss of function allele of this gene using CRISPR/Cas9 genome editing (Figure 5H and Figure S5E-F). *Or65a* mutant females completely lack responses to R-HDEA in at4 sensilla (Figure 5I), and display no preference for R-HDEA-perfumed males (Figure 5J and Figure S5D). These findings confirm OR65a is responsible for the increased female receptivity in *D. moj. wrigleyi* towards consubspecific males. We next analyzed the targeting pattern of Or65a OSNs to glomeruli in the antennal lobe (AL). Comparative morphological analyses revealed a high similarity in the basic architecture of this olfactory center in *D. moj. wrigleyi* and *D. moj. sonorensis* (Figure 6A, Movie S5-6 and File S2-3) with at least three glomeruli not present in the well-characterized AL of *D. melanogaster* (Table S1 and Figure S6A). Previous genetic tracing of at4 OSNs in *D. melanogaster* revealed projections to three glomeruli: Or47b neurons to VA1v, Or88a neurons to VA1d and Or65a/b/cneurons to DL3 (Couto et al., 2005). To label these neurons in *D. mojavensis* subspecies, we backfilled at4 sensillum neurons with a fluorescent dye. In both *D. moj*. neurons to R-HDEA (10 μg). Black-filled circles indicate significant differences from the solvent (Hexane) *wrigleyi* and *D. moj. sonorensis*, we observed projections to three glomeruli: VA1v, VA1d and a glomerulus not present in *D. melanogaster* that we named “VA8” (Figure 6B, Figure S6B and Movie 7-8, for terminology see STAR Methods). Assuming a conserved projection pattern of Or47b and Or88a neurons, we infer that the VA8 glomerulus is innervated by Or65a neurons in *D. mojavensis*.

**Figure 6.**
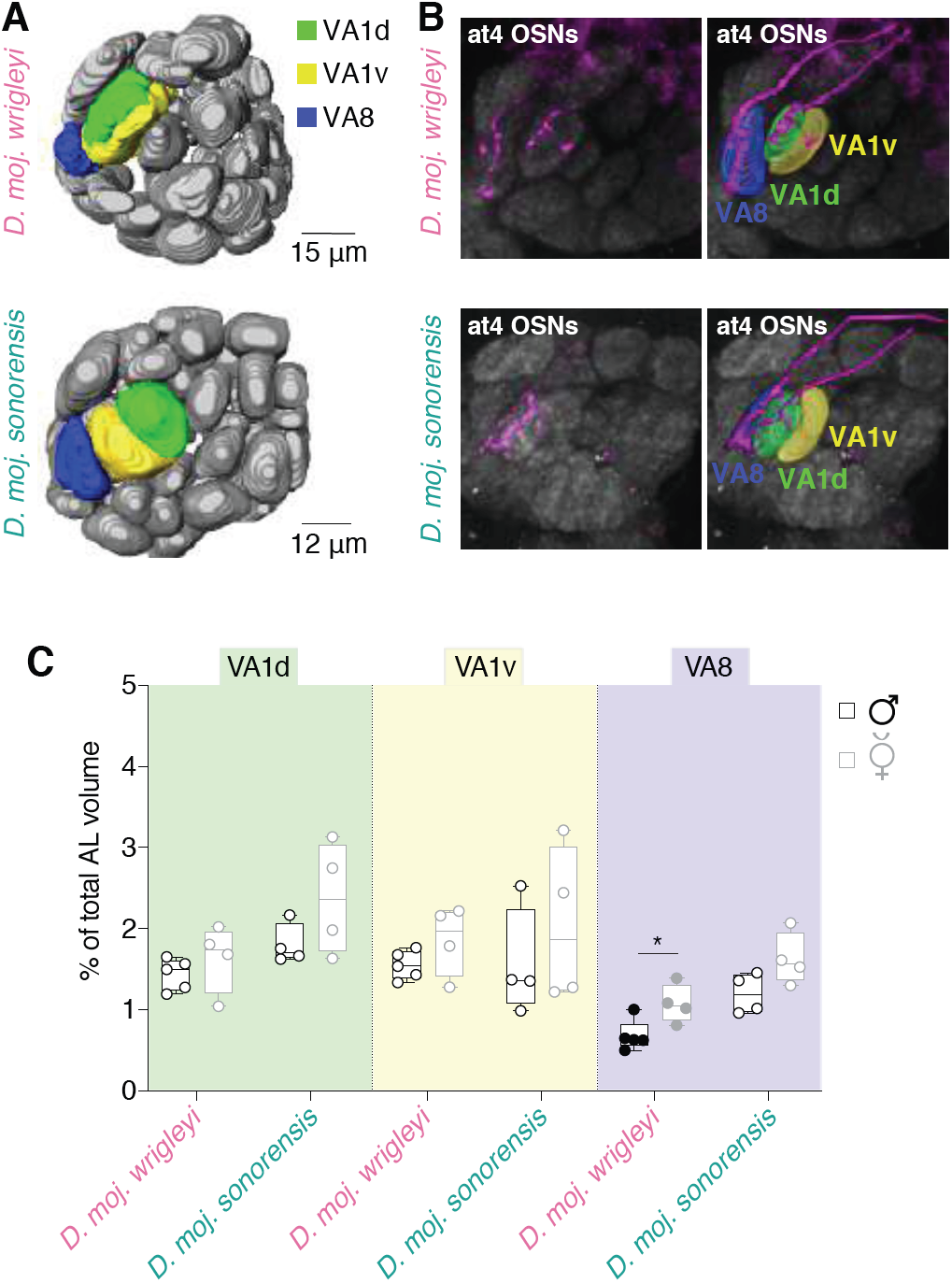
Conserved peripheral targets of R-HDEA-sensing neurons. **A**, Three-dimensional reconstruction of antennal lobes from representative female brains of *D. moj. wrigleyi* and *D. moj. sonorensis*. Neurobiotin-marked neurons in Figure 6B are highlighted: VA1d (green), VA1v (yellow), and VA8 (only present in *D. moj. wrigleyi* and *D. moj. sonorensis* in blue). VA8 is located ventrally to VA1v and anterior to VL2a in an area far off the DL3 glomerulus targeted by Or65a neurons *in D. melanogaster*. DL3 displays similar position in both *D. melanogaster* and *D. mojavensis* subspecies, see Table S1 and Figure S6C for details. Scale bar = 20 μm. **B**, Left top panel: Fluorescent staining for neurobiotin (magenta) and nc82 (grey) of *D. moj. wrigleyi* antennal lobe, backfilled from the at4 sensillum in *D. moj. wrigleyi* (identified by electrophysiological recordings; Figure 4E). Right top panel: Reconstruction of the neurobiotin-marked neurons and their corresponding glomeruli reveals at4-housed neurons project to three glomeruli (*D. mojavensis* VA8, VA1v and VA1d). Identification of glomeruli was verified by comparing the reconstructed images to the map of *D. melanogaster* AL (Grabe et al., 2015) (See STAR Methods). See Figure S6D-F for backfilling of at1 sensilla. Left panel: Neurobiotin backfilled neurons from at4 sensillum in *D. moj. sonorensis* that innervate VA8, VA1v and VA1d. Images in the four panels correspond to a projection of 40 Z-stacks (Watch Movie S7-8). See Table S1 for more details on glomerular identity and volumes in both subspecies (*D. moj. wrigleyi* and *D. moj. sonorensis*). **C**, Volumes of VA1d, VA1v, and VA8 glomeruli normalized to the total AL volume in *D. moj. wrigleyi* and *D. moj. sonorensis* (males in black, females in grey). Filled circles indicate significant difference between both sexes of the same subspecies. Mann Whitney U test, ns *P* > 0.05; * *P* < 0.05, n=4-6 brains.

Pheromone-responsive glomeruli often display sex-specific volume differences (Grabe et al., 2016; Kondoh et al., 2003). Quantitative volume analysis of the glomeruli innervated by at4 neurons in both subspecies revealed that *D. moj. wrigleyi* VA8 is the only sexually dimorphic unit with an increased volume in females (Figure 6C and Figure S6G-H). In addition, there was with a non-significant trend of larger volume of VA8 in *D. moj. sonorensis* females compared to males (Figure 6C and Figure S6G-H). Together, these results indicate that OR65a-expressing neurons in both subspecies of *D. mojavensis* detect R-HDEA and project to the same region of the AL.

### Subspecies-specific contributions of R-HDEA and auditory cues in sexual isolation

As *D. moj. wrigleyi* females can distinguish between their consubspecific males and *D. moj. sonorensis* males (which lack R-HDEA (Figure 2A)) (Figures 1C and 7A), we asked whether R-HDEA mediates this discrimination. We presented to a *D. moj. wrigleyi* female a choice of a *D. moj. wrigleyi* male perfumed with hexane and a male of *D. moj. sonorensis* perfumed with R-HDEA. Female preference for its consubspecific male was greatly reduced (Figure 7A’ and Figure S7A). Furthermore, when given a choice between two males of *D. moj. sonorensis*, of which only one was perfumed with R-HDEA, *D. moj. wrigleyi* females exhibited a strong preference for the perfumed ones (Figure 7A”).

**Figure 7.**
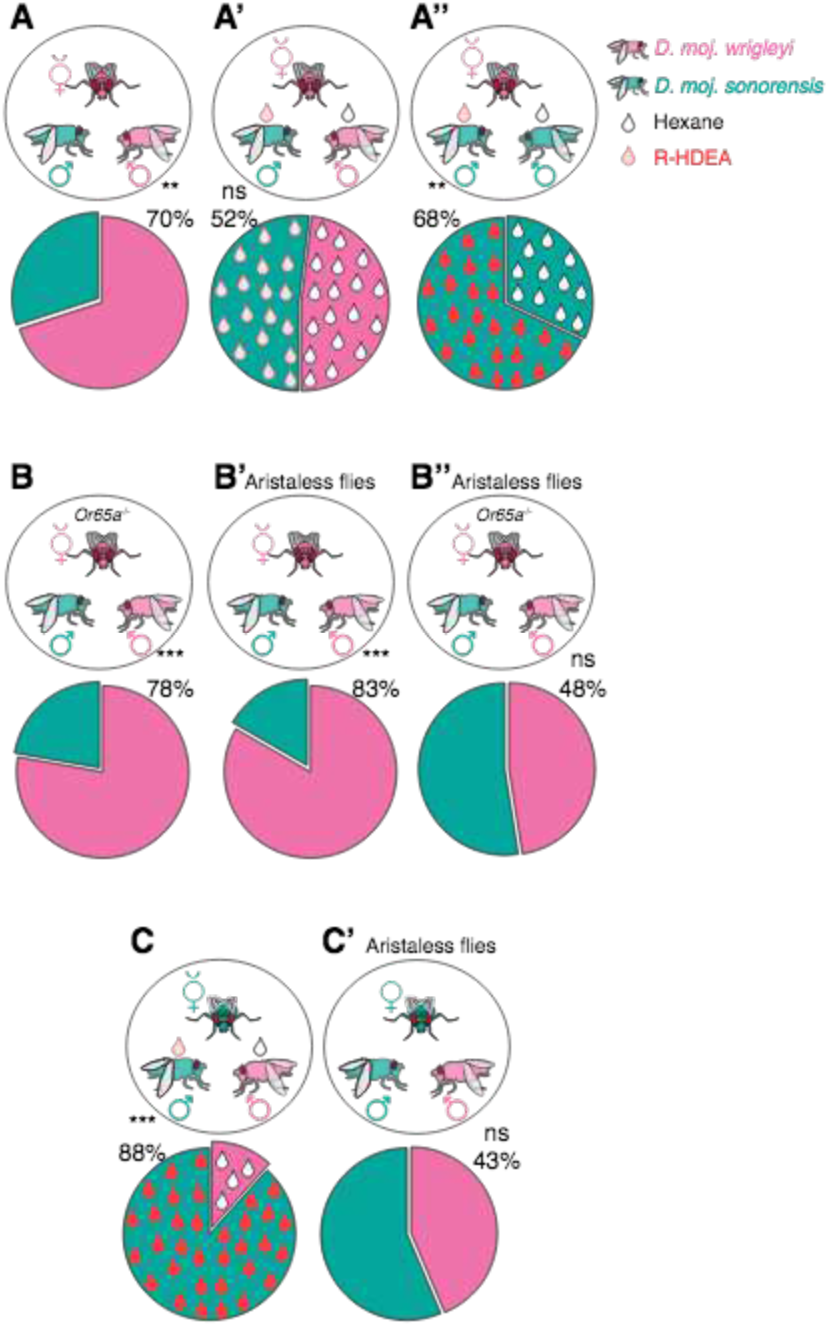
R-HDEA and auditory cues co-mediate sexual isolation among *D. mojavensis* subspecies. **A,** Competition between two males of different subspecies to copulate with a *D. moj. wrigleyi* virgin female. Pie-charts in A-C’ represent copulation success [%] of the rival males. Chi-square test, ns *P* > 0.05; ** *P* < 0.01; *** *P* < 0.001; n (A, A’, A”, B, B’, B”, C, C’) = 20, 26, 17, 18, 12, 21, 21, 22. All males and females used in this and other panels were 10-day-old virgin flies. **A’,** Competition between two males of different subspecies, *D. moj. sonorensis* perfumed with R-HDEA (red droplet) and *D. moj. wrigleyi* perfumed with hexane (black droplets), to copulate with a *D. moj. wrigleyi* virgin female. See Figure S7A for details regarding the S-HDEA and HDA-perfumed flies. **A”,** Competition between two males of *D. moj. sonorensis*, perfumed with R-HDEA (red droplet) or hexane (black droplet), to copulate with a *D. moj. wrigleyi* virgin female. **B,** Competition between two males of different subspecies, *D. moj. sonorensis* and *D. moj. wrigleyi*, to copulate with a *D. moj. wrigleyi* virgin female that lacks the R-HDEA-detecting channel OR65a. **B’,** Competition between two aristaless males of different subspecies, *D. moj. sonorensis* and *D. moj. wrigleyi*, to copulate with an aristaless *D. moj. wrigleyi* virgin female. **B”,** Competition between two aristaless males of different subspecies, *D. moj. sonorensis* and *D. moj. wrigleyi*, to copulate with an aristaless *D. moj. wrigleyi* virgin female that lacks the R-HDEA-detecting channel OR65a. **C,** Competition between two males of different subspecies, *D. moj. sonorensis* perfumed with R-HDEA (red droplet) and *D. moj. wrigleyi* perfumed with hexane (black droplets), to copulate with a *D. moj. sonorensis* virgin female. See Figure S7B for details regarding the S-HDEA and HDA-perfumed flies **C’,** Competition between two aristaless males of different subspecies, *D. moj. sonorensis* and *D. moj. wrigleyi*, to copulate with an aristaless *D. moj. sonorensis* virgin female.

Despite this instructive role of R-HDEA in mate discrimination, we observed that *D. moj. wrigleyi Or65a* mutant females (which cannot detect R-HDEA) still display strong preference for consubspecific males (Figure 7B), indicating that other cues must exist. Previous reports revealed that subspecies-specific songs during courtship induce mate recognition among *D. mojavensis* subspecies (Etges et al., 2007; Etges et al., 2006). We therefore investigated whether auditory inputs are sufficient to mediate mate recognition. Aristaless *D. moj. wrigleyi* females – lacking an essential part of the antennal auditory organ – still exhibited normal preference to aristaless consubspecific males over aristaless hetersubpecific ones (Figure 7B’). However, aristaless, *Or65a* mutant females exhibited a major loss in the recognition of consubspecific males (Figure 7B”). These results indicate that R-HDEA and auditory cues have redundant roles in permitting subspecies discrimination of *D. moj. wrigleyi*.

In contrast to *D. moj. wrigleyi* females, *D. moj. sonorensis* females preferred consubspecific males over the *D. moj. wrigleyi* one, even when the consubspecific one was perfumed with R-HDEA or other male-specific compounds (Figure 7C and Figure S7B). This subspecies appears to rely solely on auditory input for subspecies recognition, as aristaless females displayed indiscriminate preference for consubspecific and hetersubpecific males (Figure 7C’).

## Discussion

Modifications in sex pheromones and their cognate sensory detection systems have been proposed as the most rapid coevolving elements permitting mate recognition and speciation (Smadja and Butlin, 2009). Our results provide numerous novel insights to this co-evolutionary framework, all of which represent, in concert, intra- and interspecific sexual barriers.

First, among *D. mojavensis* subspecies, we identified four male-specific acetates of which three were exclusively produced by males of the northern subspecies (Figure 2A). This dramatic change in the males profiles is surprising given the relatively short divergence time between *D. mojavensis* subspecies (Etges, 2019; Matzkin, 2014). However, although these subspecies utilize different host cacti in nature (Matzkin, 2014), in our study they were bred on the same food suggesting that the different chemical profiles are not a consequence of nutrition and genetically inherited. The male-specificity and sexual transfer of these acetates resembles *c*VA in *D. melanogaster*, which is produced by the ejaculatory bulb and transferred to females during mating. Our data reveals that the opposite *c*VA-promoted behaviors among sexes (Kurtovic et al., 2007; Zawistowski and Richmond, 1986) – inhibiting male and inducing female mating behaviors – are mediated by two different compounds in *D. mojavensis* (Figures 3B and 4B). While R-HDEA induces female receptivity in the subspecies producing it, OCDA is present and suppresses male courtship in all *D. mojavensis* subspecies. Consistent with OCDA-induced behaviors, a recent study (Chin et al., 2014) showed that *D. mojavensis* males avoid courting females perfumed with ejaculatory-bulb extracts of mature males.

Second, *D. mojavensis* subspecies display identical courtship elements, indicating that the sexual isolation (Figure 1C) is due to different sexual traits and preferences. Our results address a longstanding mystery of the modalities that mediate prezygotic isolation between *D. mojavensis* subspecies. Through a combination of behavioral competition experiments, genome engineering, and arista removal (Figure 7), we reinforced that the dedicated R-HDEA-sensing neurons and auditory cues are co-mediating mate recognition in *D. moj. wrigleyi*. In contrast, R-HDEA did not induce any behavioral change in *D. moj. sonorensis* females that rely only on auditory cues (i.e., the subspecies-specific song (Etges et al., 2006)) for mate recognition. The striking change of R-HDEA-induced behaviors (Figure 4C and Figure S4D), despite its conserved peripheral sensory pathway (Figure 5E and 6B-C), implies upstream modulations in the processing pathways among *D. mojavensis* subspecies. These findings are concordant with dissimilar cVA-promoted behaviors among sexes of *D. melanogaster* (Kohl et al., 2015), likely because of dimorphic wiring in the central brain (Kohl et al., 2013; Ruta et al., 2010) in both sexes.

Third, species of the *Drosophila* genus show species-specific courtship behaviors (Spieth, 1952), which might include nuptial gift donation (Steele, 1986), partners’ song duet (LaRue et al., 2015; O’Grady and Markow, 2012) or territorial dating (O’Grady and Markow, 2012). To our knowledge, although males of *D. melanogaster* and some Hawaiian *Drosophila* species have been shown to attract females over distance by releasing pheromones in fecal droplets (Mercier et al., 2018; O’Grady and Markow, 2012), releasing droplets during close courtship has not been described for drosophilid flies yet. We demonstrate that *D. mojavensis* subspecies males advertise their presence by producing a droplet of anal secretions in close proximity to females (Figure S1). In *D. moj. wrigleyi*, this droplet contains the volatile sex pheromone R-HDEA (Figure S2G) that increases female receptivity. Such a trait might be supportive to the observed sexual behavior of *D. mojavensis* in nature, where males occupy undamaged areas next to the necrotic feeding sites on cacti and attract females to this spot (O’Grady and Markow, 2012).

Fourth, chemical analyses revealed absence of *c*VA in *D. mojavensis* that instead possesses other novel analogous stimuli. Such evolutionary change raises the possibility of a prompt divergence in the tuning of the corresponding sensory receptors (Crowley-Gall et al., 2016; Linz et al., 2013; Prieto-Godino et al., 2017). Here, we demonstrate a distinct modification in the peripheral perception of *D. mojavensis*, compared to *D. melanogaster*, of the newly-identified pheromones. Accordingly, we demonstrate that the diverged OR65a orthologs of both species (Figure S4J) have different functional properties; the *D. melanogaster-Or65a* is not detecting the best ligand of *D. moj. wrigleyi* OR65a, R-HDEA, and *vice versa* (Figure 4E and Figure S4I). In line with the functional divergence, the number of Or65 copies is frequently changing along the *Drosophila* phylogeny (Guo and Kim, 2007), indicating that the variation at this locus could be important for the evolution of novel olfactory channels (Guo and Kim, 2007).

Finally, divergence of the chemosensory genes among the closely-related species could be accompanied by a physiological alteration in the underlying central circuitry (Seeholzer et al., 2018). Our results provide, to our knowledge, the first correlation between anatomical difference and divergent behaviors of homologous neurons. Or65a neurons that in *D. melanogaster* and *D. mojavensis* maintain orthologous receptor expression (*Or65a*) and sensillum identity (at4) have evolved novel projection patterns in the AL, with Or65a targeting DL3 in *D. melanogaster* and VA8 in *D. mojavensis*. The intriguing wiring of the Or65a in *D. mojavensis* indicates that it could be partnering with another subset of second-order neurons responsible for the different type of sexual behaviors: Or65a mediates female receptivity in *D. moj. wrigleyi*, whereas it inhibits female attraction toward cVA in *D. melanogaster* (Lebreton et al., 2014).

Future evolutionary investigations of the Or65a innervation pattern in the higher brain centers will shed the light on the divergent functions of R-HDEA among the closely related subspecies of *D. mojavensis* and will explain how neural circuits coevolve with sex pheromones to permit specific mate recognition.

## Acknowledgements

We thank R. Stieber for performing immunohistochemistry experiments, I. Alali for fly rearing and maintenance, L. Zhang for the preliminary molecular data, and J. Layne for his technical help in confocal imaging. We are grateful to K. Weniger, Sybille Lorenz, S. Trautheim, C. Hoyer and L. Merkel for their technical support, and to J.R. Arguello, L. Prieto-Godino, W.B. Walker III and members of the Department of Evolutionary Neuroethology, MPI-CE, in Jena, Germany for discussions. Transgenic flies were obtained from the Bloomington *Drosophila* Stock Center (NIH P40OD018537), and wild-type flies were obtained from the San Diego *Drosophila* Species Stock Center (now The National *Drosophila* Species Stock Center, Cornell University). This research was supported through funding by the Max Planck Society. T.O.A. is supported by a Human Frontier Science Program Long-Term Fellowship (LT000461/2015-L). B.A. is supported by a National Science Foundation grant (IOS-1456932). D.Z and C.L are supported by the Royal Physiographic Society of Lund and the Swedish Research Council. L.M.M. lab was partly supported by a National Science Foundation grant (IOS-1557697). Research in R.B.’s laboratory is supported by ERC Consolidator and Advanced Grants (615094 and 833548, respectively).

## Author Contributions

M.A.K, H.K.M.D., B.S.H., and M.K. conceived of the project. All authors contributed to experimental design, analysis and interpretation of results. M.A.K. prepared all figures. M.A.K. with the help of T.O.A. designed the genetic constructs and T.O.A. performed the microinjections for new drosophilid mutants. S.L generated reagents for the transgenic flies. A.S. identify and J.W. synthesized the novel compounds. Other experimental contributions were as follows: M.A.K (Fig.1, Fig.2, Fig. 3, Fig. 4, Fig. 5C and E-J, Fig. 6B, Fig. 7, Fig. S1, Fig. S2A-C and F-G, Fig. S3, Fig. S4, Fig. S5A and C-G, Fig. S6B, D-F, Fig. S7, Movie S1-4 and 7-9, and File S1), T.O.A (Fig. 5H), V.G and A.D (Fig. 6, Fig. S6A-E and F-G, Movie 5-9 and Table S1), B.A. (Fig. 5A-B), D.Z. and C.L. (Fig. 5D and Fig. S5B), F.K. (Fig. S2B-C), L.M.M. (Fig. 1B and Fig. S6D). M.A.K. wrote the original manuscript, and M.K., T.O.A, R.B. and B.S.H. contributed to the final manuscript. All coauthors contributed to the subsequent revisions.

## STAR Methods

### *Drosophila* lines and chemicals

#### Fly stocks

Wild-type flies used in this study were obtained from the *National Drosophila* Species Stock Centre (NDSSC) (https://stockcenter.ucsd.edu). Stock numbers and breeding diets are listed in Key Resources Table. Transgenic *D. melanogaster* flies and *D. mojavensis* knock-out lines generated in this study are listed in Key Resources Table. All flies were reared at 25 °C, 12h Light: 12h Dark and 50% relative humidity. For more details on the food recipes see *Drosophila* Species Stock Centre (http://blogs.cornell.edu/drosophila/recipes/). Animals subjected to arista or foreleg removal were left to recover for 2 days post-surgery before behavioral experiments.

#### Chemicals

Odorants used in this study, their source, and their CAS numbers are listed in the Key Resources Table. All odors were diluted in dichloromethane (DCM) for single sensillum recordings (SSR), in dimethyl sulfoxide (DMSO) for oocytes and tip recording experiments or in hexane for behavioral experiments.

### Chemical analyses

#### Thermal Desorption-Gas Chromatography-Mass Spectrometry (TD-GC-MS)

Individual flies or ejaculatory bulbs were prepared or dissected, respectively, for chemical profile collection as described previously (Dweck et al., 2015) with some modifications. Briefly, the GC-MS device (Agilent GC 7890A fitted with an MS 5975C inert XL MSD unit; www.agilent.com) was equipped with an HP5-MS UI column (19091S-433UI; Agilent Technologies). After desorption at 250 °C for 3 min, the volatiles were trapped at −50 °C using liquid nitrogen for cooling. In order to transfer the components to the GC column, the vaporizer injector was heated gradually to 270 °C (12 °C/s) and held for 5 min. The temperature of the GC oven was held at 50 °C for 3 min, gradually increased (15 °C/min) to 250 °C and held for 3 min, and then to 280 °C (20 °C/min) and held for 30 min. For MS, the transfer line, source and quad were held at 260 °C, 230 °C and 150 °C, respectively. Eluted compounds for this and the following analyses were ionized in electron ionization (EI) source using electron beam operating at 70 eV energy and their mass spectra were recorded in positive ion mode in the range from *m/z* 33 to 500. Anal-droplets of courting males were collected on cover glass (22 × 22 mm, Cat. No.: 631-0126, https://uk.vwr.com) and then analyzed by TD-GC-MS. The structures of the newly identified acetates were confirmed by comparing their mass spectrum with synthesized standards.

#### Hexane-body washes analysis by GC-MS

Fly body extracts were obtained by washing single flies of the respective sex, age and mating status in 10 μl of hexane for 30 min. For GC stimulation, 1 μl of the odor sample was injected in a DB5 column (Agilent Technologies; www.agilent.com), fitted in an Agilent 6890 gas chromatograph, and operated as described previously (Stokl et al., 2010). The inlet temperature was set to 250 °C. The temperature of the GC oven was held at 50 °C for 2 min, increased gradually (15 °C/min) to 250 °C, which was held for 3 min, and then to 280 °C (20 °C/min) and held for 30 min. The MS transfer-line, source and quad were held at 280 °C, 230 °C and 150 °C, respectively. Average amounts of HDEA, HDA and OCDA from individual flies were quantified by comparing their peak areas with the area of hexadecane (10 ng), which was added to the fly extract as an internal standard.

#### Chiral chromatography

To check the ratio of (2*R*,10Z)-10-Heptadecen-2-yl acetate and (2*S*,10*Z*)-10-Heptadecen-2-yl acetate (R/S-HDEA), hexane body extracts of male flies were injected into a CycloSil B column (112-6632, Agilent Technologies; www.agilent.com) fitted in Agilent 6890 gas chromatograph and operated as follows: The temperature of the GC oven was held at 40 °C for 2 min and then increased gradually (10 °C/min) to 170 °C, then to 200 °C (1 °C/min), and finally to 230 °C (15 °C/min) which was held for 3 min.

#### Matrix-assisted laser desorption/ionization-time of flight (MALDI-TOF)

MALDI-TOF experiments of 13-day-old flies were performed on MALDI Micro MX (Waters, UK) operated in a reflectron mode with acceleration and plate voltages at 12 and 5 kV, respectively. Due to the relative high volatility and weaker ionization of R&S-HDEA and HDA compared to OCDA, only OCDA signals were detected in the MSI spectra. Moreover, OCDA was predominantly present in a form of [M+K]^+^ adducts. Delayed extraction time was 500 ns. Compound’s desorption/ionization was realized by nitrogen UV laser (337 nm, 4 ns pulse of maximum 320 μJ and frequency of 20 Hz). Matrix ions were suppressed with a low mass cut-off set at *m/z* 150. Samples of the flies were fixed on their backs and processed as described previously (Kaftan et al., 2014). The number of laser shots per spot was optimized and set to 60 (128 μJ/shot). The range of the measured masses was set from *m/z* 100 to *m/z* 1000. Data were collected with MassLynx 4.0 software and processed with custom-made software MALDI Image Convertor (Waters, UK) to obtain spatially differentiated data. These data were exported to the BioMap software (Novartis, Switzerland) and converted to 2-D ion intensity heat maps. All samples were analyzed in positive ion mode and imaged using a step size of 100 μm. Methanolic solution of LiDHB and LiVA (in case of copulated flies) matrix in a concentration of 20 mg/mL was sprayed on the fly samples by an airbrush. For one sample, 0.4 mL of LiDHB/LiVA matrix solution was used to form approximately 25 layers. Waiting time between two consecutive sprays was 3s. Lithium 2,5-dihydroxybenzoate (LiDHB) and lithium vanillate (LiVA) matrix were synthesized as described previously (Cvacka and Svatos, 2003; Horka et al., 2014). Polyethylene glycol oligomers (with an average molecular weight of 200, 300, 600 and 1000 Da) for calibration of the mass spectrometer were purchased from Sigma-Aldrich as well as precursors of the LiDHB and LiVA. Potassiated adducts of OCDA at *m/z* 487 evinced a negative mass shift which was observed within a range from *m/z* 0.05 to *m/z* 0.3 between measurements.

#### Fly odor analysis by Solid Phase Microextraction (SPME)

Volatiles were collected from 20 *D. moj. wrigleyi* males trapped in a mesh inside a capped 4 ml glass vials with polytetrafluoroethylene-lined silicone septa (Sigma, 23242-U) to preclude flies contact to the SPME fiber (grey hub plain; coated with 50/30 μm divinylbenzene/carboxen on polydimethylsiloxane on a StableFlex fiber, Sigma, 57328-U). SPME fiber was exposed to the trapped flies for 1 h at room temperature. The SPME fiber was then retracted and immediately inserted into GC-MS system (Agilent 7890B fitted with MS 5977A unit) as operated previously (Date et al., 2013). The inlet temperature was set to 250 °C. The temperature of the GC oven was held at 40 °C for 3 min and then increased gradually (5 °C/min) to 280 °C, which was held for 10 min.

#### Perfuming flies with hexane or male-specific compounds

Males and female flies were perfumed with the acetates singly diluted in hexane or hexane alone as previously described (Dweck et al., 2015). Briefly, each fly was coated by ~ 1-3 ng of the compound of interest after evaporating the hexane under nitrogen gas flow. Flies were transferred to fresh food to recover for 2 h and then introduced to the courtship arenas or subjected to GC-MS analysis to confirm the increased amount of the perfumed acetate.

##### Chemical identification and synthesis

(provided as a separate word file)

### Behavioral experiments

#### Single and competitive mating assays

Males and females were collected after eclosion and raised individually and in groups, respectively. For each experiment, courtship assays were performed in a (1 cm diameter × 0.5 cm depth) chamber covered with a plastic slide. Courtship behaviors were recorded for 10 or 20 min using a GoPro Camera 4 or Logitech C615 as stated in the figure legends. All single mating experiments were performed under red light (660 nm wavelength) at 25 °C and 70% humidity. Each video was analyzed manually for copulation success, which was measured by the percentage of males that copulated successfully in the first 10 min, copulation latency, which was measured as the time taken by each male until the onset of copulation, and courtship index, which was calculated as the percentage of time that the male spends courting the female during 10 min. In Figure 2B, copulation success experiments were manually monitored for 1 h. Freeze-killed females were used in the courtship assays to disentangle male sexual behaviors from female acceptance.

In the competition mating assays, rival flies were marked by UV-florescent powder of different colors (red: UVXPBR; yellow: UVXPBB; green: UVXPBY; purchased from Maxmax.com; https://maxmax.com/) 24 hours before the experiments. Competition assays were manually observed for 1 h and copulation success was scored identifying the successful rival under UV light. Females killed by freezing were used to calculate the courtship index of males in presence of the different acetates or hexane perfumed on the females’ body. For tarsi- or arista-less flies either the first three segments of male tarsi or both aristae were clipped with a clean razor blade and flies were kept to recover for two days on fresh food before introduction into the courtship arena. All courtship and copulation data were acquired by a researcher blinded to the treatment. For collecting male anal droplets, a virgin male and a decapitated female were kept in a courtship chamber covered with a glass coverslip. Males were allowed to court and discharge the anal-droplets on the glass coverslips which were then cracked to small pieces and inserted into the TD-GC-MS tubes for analysis. As a control, glass coverslips were sampled from courtship arenas containing only males.

#### Wind tunnel assay

Long-range attraction experiments were performed in a wind tunnel as described previously (Becher et al., 2010). The wind tunnel was maintained within a climate chamber at 25 °C and 50-55% humidity in white light. Flies were starved for approximately 20 h. Per assay, 5 females or males (10-15 days old) were released in presence of a filter paper (3 × 3 cm) charged with either 100 μl of cactus juice (Opuntia, https://www.luckyvitamin.com) alone or with 10 μl of acetates singly or in a mix. Flies landing on the filter paper were counted for the first 10 min after release. **T-maze assay.** Short-range attraction experiments were carried out as described in (Dweck et al., 2016). In brief, 30 starved females and males (10-15 days old) were allowed to choose between two sides containing 0.5 ml agar (1%) charged with either 50 μl of cactus juice alone or the different male-specific acetates. The attraction index (AI) was calculated as in (Dweck et al., 2016). Experiments were carried out in a climate chamber at 25 °C and 50% humidity. All T-maze assays were performed under white light.

### Electrophysiological and molecular biology experiments

#### Single sensillum recording (SSR)

Adult flies were immobilized in pipette tips, and the third antennal segment was placed in a stable position onto a glass coverslip (Olsson and Hansson, 2013). Sensilla types were localized under a microscope (BX51WI; Olympus) at 100x magnification and identified by diagnostic odors stated in Key Resources Table. The extracellular signals originating from the OSNs were measured by inserting a tungsten wire electrode in the base of a sensillum and a reference electrode into the eye. Signals were amplified (Syntech Universal AC/DC Probe; Syntech), sampled (10,667.0 samples/s), and filtered (300 – 3,000 Hz with 50/60 Hz suppression) via USB-IDAC connection to a computer (Syntech). Action potentials were extracted using AutoSpike software, version 3.7 (Syntech). Synthetic compounds were diluted in dichloromethane, DCM, (Sigma-Aldrich, Steinheim, Germany). Prior to each experiment, 10 μl of diluted odors were freshly loaded onto a small piece of filter paper (1 cm^2^, Whatman, Dassel, Germany), and placed inside a glass Pasteur pipette. The odorant was delivered by placing the tip of the pipette 2 cm away from the antennae. Neuron activities were recorded for 10 s, starting 2 s before a stimulation period of 0.5 s. Responses from individual neurons were calculated as the increase (or decrease) in the action potential frequency (spikes/s) relative to the pre-stimulus frequency. Traces were analyzed by sorting spike amplitudes in AutoSpike and then analyzed in Excel and processed in Adobe Illustrator CS (Adobe systems, San Jose, CA).

#### Tip recording

Tip recordings were performed as described previously (Moon et al., 2006) from tarsal sensilla. Male flies (8-10 day old) were immobilized in pipette tips, and the foreleg was fixed with Scotch tape onto a glass coverslip. A reference glass electrode filled with Ringer’s solution (140 mM NaCl, 3 mM MgCl2, 2 mM CaCl2, 10 mM D-glucose, 10 mM HEPES, pH 7.4) was inserted into the thorax of the fly. The different tarsal sensilla were stimulated by placing a glass capillary filled with OCDA (10 μg) dissolved in DMSO on the sensillum tip. The recording electrode was connected to a pre-amplifier (TastePROBE, Syntech, Hilversum, The Netherlands), and the signals were collected and amplified (10x) by using a signal-connection interface box (Syntech) in conjunction with a 100-3000 Hz band-pass filter. Action potentials measurements were acquired with a 9.6 kHz sampling rate and analyzed with AutoSpike.

#### RNA extraction, cDNA synthesis

Total RNA from single flies of *D. mojavensis* subspecies and from 100 antennae was extracted using an RNA isolation kit (Directzol^TM^ RNA MiniPrep, Zymo Research). First strand cDNA was generated from 1.0 μg of total RNA, using oligo-dT20 primers and Superscript™ III (Thermo Fisher Scientific). Derived cDNAs were used to amplify housekeeping and olfactory receptor ORFs via PCR with MyTaq™ DNA Polymerases (Bioline) and primers listed in Key Resources Table. PCR amplicons were cloned into pCR4-TOPO (Invitrogen) and verified by sequencing.

#### Sequence alignment

Available (flybase, https://flybase.org) and generated (in our study) protein-coding regions of *Or65a* and *Or47b* were analysed in Geneious (v11.0.5). Briefly, a multiple sequence alignment was generated using the MAFFT (v7.309) tool with E-INS-I parameters and scoring matrix 200 PAM / K=2 as previously described (Katoh and Standley, 2013). The final tree of OR65a orthologs was reconstructed using a maximum likelihood approach with the GTR+G+I model of nucleotide substitution and 1000 rate categories of sites in Fasttree (v2.1.5). The tree was visualized and processed in Geneious.

#### Functional analysis of receptor genes in *Xenopus* oocytes

Oocyte injections and two-electrode voltage clamp recordings were described previously (Zhang and Lofstedt, 2013). Briefly, the open reading frames of *D. moj. wrigelyi Or47bl, Or47b2, Or65a, Or67d, Or88a* and *O*rco were amplified from *D. moj. wrigelyi* cDNA using primers (Key Resources Table) adding BamHI (fwd) and Xbal (rev) restriction sites, and a Kozak sequence (GCCACC) immediately upstream of the first ATG. The PCR products were digested with BamHI and Xbal and sub-cloned into the expression vector pCS2+. Maxi preps of recombinant plasmids were linearized with NotI and transcribed to cRNAs with mMESSAGE mMACHINE SP6 kit (Thermo Fisher Scientific, Waltham, MA, USA). *X. laevis* (purchased from Xenopus Express France, Vernassal, Haute-Loire, France) oocytes were defolliculated with 1.5 mg/mL collagenase (Sigma-Aldrich Co., St. Louis, MO, USA) in Oocyte Ringer 2 solution (82.5 mM NaCl, 2 mM KCl, 1 mM MgC12, 5 mM HEPES, pH 7.5). cRNAs of each OR together with *D. moj. wrigelyi* ORCO cRNA (50 ng each) were co-injected into the oocytes and incubated at 18 °C for 3-5 days prior to recordings. Stock solutions of the tested compounds were prepared in dimethyl sulfoxide (DMSO) (Sigma-Aldrich Co., St. Louis, MO, USA) and diluted to the indicated concentrations with Ringer’s buffer (96 mM NaCl, 2 mM KCl, 5 mM MgC12, 0.8 mM CaCl2, 5 mM HEPES, pH 7.6). The compounds mentioned in Key Resources Table were applied to the oocytes successively at a rate of 2 ml/min with extensive washings by Ringer’s buffer in the intervals. Whole-cell inward currents were recorded by two-electrode voltage clamp with a TEC-03BF amplifier (npi electronic GmbH, Tamm, Germany) at the holding potential of −80 mV. Data were collected and analyzed by Cellworks software (npi electronic GmbH, Tamm, Germany).

#### Expression of olfactory receptors in *D. melanogaster* at1 neurons

Transgenic lines were generated according to standard procedures as described (Gonzalez et al., 2016). The open reading frames of *D. moj. wrigelyi Or47bl, Or47b2, Or65a, Or67d, Or88a* and *D. moj. sonorensis Or65a* receptors were subcloned from the corresponding pCS2+ constructs (see above) via digestion with BamHI (Cat.# R0136, New England Biolabs) and Xbal (Cat.#R0145 New England Biolabs) and ligated into pUASt.attb (Bischof et al., 2007) (a gift from Dr. Johannes Bischof) digested with BglII (Cat.#R0144, New Englands Biolabs) and Xbal (Cat.#R0145, New England Biolabs). Homozygous *UAS-OrX* lines (with transgene insertions into chromosome II) were generated at Bestgene (https://www.thebestgene.com). An OR67d-knock-out/Gal4-knock-in stock (provided by Dr. Barry J. Dickson) was individually crossed to each of the transgenic *UAS-DmojORx* flies, and homozygous lines expressing the Or gene of interest in the decoder at1 neuron of *D. melanogaster* were established. Each UAS-transgenic line was confirmed by sequencing of genomic DNA prepared from the final stocks.

### Generation of knock-out flies

#### *Drosophila* microinjections

Transgenesis of *D. mojavensis wrigleyi* was performed in-house following standard protocols (http://gompel.org/methods). For egg-laying agar plates a few g of Formula 4-24^®^ Instant *Drosophila* Medium, Blue (Carolina Biological Supply Company) soaked in water were added on the surface. Embryos were manually selected for the appropriate developmental stage prior to alignment and injection. For CRISPR/Cas9-mediated genome engineering a mix of two sgRNAs (see Key Resources Table for sequences) targeting the *white* locus, two sgRNAs targeting the *Or65a* locus (3 μM each) and Cas9 protein (2 μM) (all Synthego; sgRNAs with 2’-O-methyl 3’ phosphorothioate modifications in the first and last 3 nucleotides) was prepared. Prior to injection individual sgRNAs were mixed with Cas9 protein (1.5:1) and incubated at room temperature for 10 minutes. All concentrations are the final values in the injection mix.

#### Genotyping

Individual G0 flies were crossed to wildtype flies and G1 adults visually screened for white-eyed flies. Single white-eyed flies were subsequently crossed to wildtype flies and G2 flies genotyped by PCR. Genomic DNA of a single wing of each fly was isolated using the MyTaq™ Extract-PCR Kit (Bioline Cat No: BIO-21126), the sgRNA-target site PCR amplified (see Key Resources Table for genotyping primers) and the amplicon sanger-sequenced. Sequencing results were compared to the reference sequence using TIDE (Brinkman et al., 2014) to deconvolute sequencing traces. G2 flies, which displayed heterozygous loss-of-function mutations at the *Or65a* or white locus, were incrossed and stocks of *Or65a* and *white* loss-of-function alleles established.

#### Labeling and antennal lobe reconstruction

##### *In-situ* hybridizations

Whole-mount single *in-situ* hybridization was performed as previously described (Crowley-Gall et al., 2016; Saina and Benton, 2013). In brief, both sense and antisense digoxigenin (DIG) RNA probes were generated for each odorant receptor gene using a DIG-RNA labeling kit (Roche, Indianapolis, IN, USA) according to manufacturer’s instructions. See Key Resources Table for details about the oligonucleotides’ sequences. RNA probes were hydrolysed (60 mM Na_2_CO_3_, 40 mM NaHCO_3_, pH 10.2) at 60 °C for 1 h, ethanol precipitated and stored in formamide at −80 °C until use. Heads of *D. moj. wrigleyi* and *D. moj. sonorensis* females were cold fixed (4% paraformaldehyde, 0.05% Tween 20 in 1x PBS) for 1 h and then washed 3x 10 min in PBST (1x PBS, 0.1% Tween 20). Third antennal segments were dissected into cold fix, washed 3x 10 min in PBX (1x PBS, 1% Triton-X) and incubated for 2 h in hybridization (Hyb) buffer (50% formamide, 5x SSC, 0.05 mg/ml heparin, 0.1% Tween 20). Antennae were hybridized overnight at 55 °C using appropriate RNA probe(s) and the next day washed 5x 2 h in Hyb buffer at 55 °C, with the last wash leading to overnight incubation at 55 °C. A wash of 20 min in Hyb buffer at 55 °C, and 3x 10 min wash in PBST at room temperature were subsequently performed. Antennae were incubated in 1:500 anti-DIG-POD (in PBST and 1x BSA)for3 h followed by 3x 10 min wash in PBST. Samples were then incubated in TSA-Plus Cyanine 5 (Cy5) according to manufacturer instruction (Perkin Elmer, Waltham, MA, USA) for 70 min followed by 5x 10 min washes in PBST. Samples were subsequently suspended in 80% glycerol for visualization using confocal microscopy. Confocal Z-stacks were acquired using a Nikon A1Rsi inverted confocal microscope. ORNs were counted using NIS Elements Viewer (Melville, NY, USA).

##### Antennal lobe reconstruction

Fly brain dissections and stainings were performed as described previously (Dekker et al., 2006; Grabe et al., 2016) with some modifications as follow: Brains were dissected in PBS and fixed in 4% paraformaldehyde (PFA) for 30 min at room temperature (RT), rinsed 3x 15 min in PBS with 0.3% Triton X-100 (PT), followed by incubation with mouse monoclonal nc82 antibody (1:30, CiteAb, A1Z7V1) in 4% normal goat serum (NGS,) in PT (48 h at 4 °C). Samples were washed 4x 20 min in PT, incubated overnight with Alexa633-conjugated anti-mouse antibody (1:250, A21052, Invitrogen) in NGS-PT, rinsed 4x 20 min in PT and mounted in Vectashield (Vector Laboratories). Images were acquired with a Zeiss 710 NLO Confocal microscope using a 40x or 63x water immersion objective. Reconstruction of whole ALs and of individual glomeruli (4-6 AL specimens for each sex/species) was performed manually using the segmentation software AMIRA version 5.6.0 (FEI Visualization Sciences Group). Identification of glomeruli was verified by comparing the reconstructed images to the map of *D. melanogaster* AL (Grabe et al., 2015). Glomerular volume was calculated from reconstructed glomeruli and using the information on the voxel size from the laser scanning microscopy scans.

##### OSN backfilling

Trichoid sensilla were identified by SSR using diagnostic odor-evoked responses to R-HDEA for at4 or cVA for at1. After identifying the right sensillum, the recording electrode was removed and neurons were backfilled by placing the sensillum inside a glass capillary filled with neurobiotin (Invitrogen, 2% w/v in 0.25 M KCl). Neurobiotin was allowed to diffuse into the OSNs for 2 h. Brain staining was performed as described above, and neurobiotin was visualized using streptavidin conjugated to Alexa Fluor 555 (1:500, S32355, Invitrogen).

Due to its position we name the novel glomerulus innervated by at4 neurons in *D. mojavensis* VA8. VA8 is located ventrally to VA1v and anterior to VL2a in an area far off the DL3 glomerulus normally targeted by at4 neurons *in D. melanogaster*.

##### Scanning Electron Microscopy

Antennal scans were performed as previously described (Grabe et al., 2016). Images of the third antennal segment were acquired at 4.5k x magnification using a LEO 1450 VP scanning electron microscope with 10 kV and 11 mm working distance (Carl Zeiss).

#### Phylogenetic analysis, statistics and figure preparation

##### Phylogenetic analysis

The phylogenetic relationship among the four *D. mojavensis* subspecies and *D. melanogaster* was determined using the recent genomic analysis of *D. mojavensis* (Allan and Matzkin, 2019) and the *D. melanogaster* FlyBase assembly (FB2019_04). The BUSCO application (Waterhouse et al., 2018) was ran using the Diptera dataset (composed of 2,799 loci) for all five genomes and a common set of 2,622 loci was obtained. Individual gene alignments using MUSCLE (Edgar, 2004) were then concatenated and 3^rd^ base nucleotide positions (1,887,113 sites) were extracted. This dataset was subsequently ran on PhyML (Guindon et al., 2010) under a GTR +G+I model with 1,000 bootstraps to assess the support for the phylogenetic inference. For the receptors’ phylogeny, a multiple sequence alignment was generated using the MAFFT (v7.309) tool with E-INS-I parameters and scoring matrix 200 PAM / K=2 (Katoh and Standley, 2013). The final tree was reconstructed using a maximum likelihood approach with the GTR+G+I model of nucleotide substitution and 1000 rate categories of sites in Fasttree (v2.1.5). The tree was visualized and processed in Geneious (v11.0.5).

##### Statistics and figure preparation

Normality was first assessed on datasets using a Shapiro test. Statistical analyses (see the corresponding legends of each figure) and preliminary figures were conducted using GraphPad Prism v. 8 (https://www.graphpad.com). Figures were then processed with Adobe Illustrator CS5.

#### Data and biological material availability

All relevant data supporting the findings of this study and all unique biological materials generated in this study (e.g., mutant and transgenic fly strains) are available from the corresponding authors upon request.

---

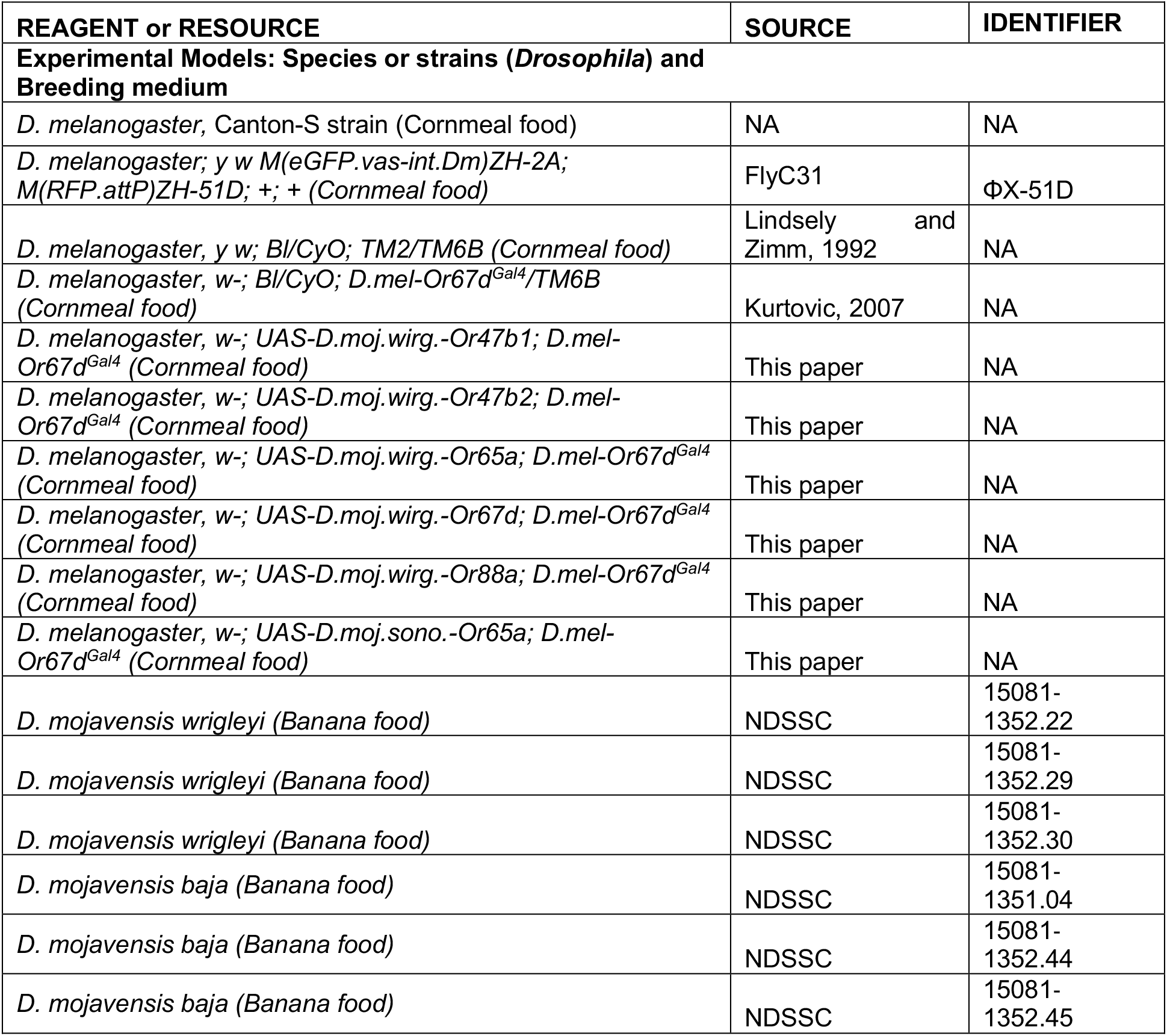
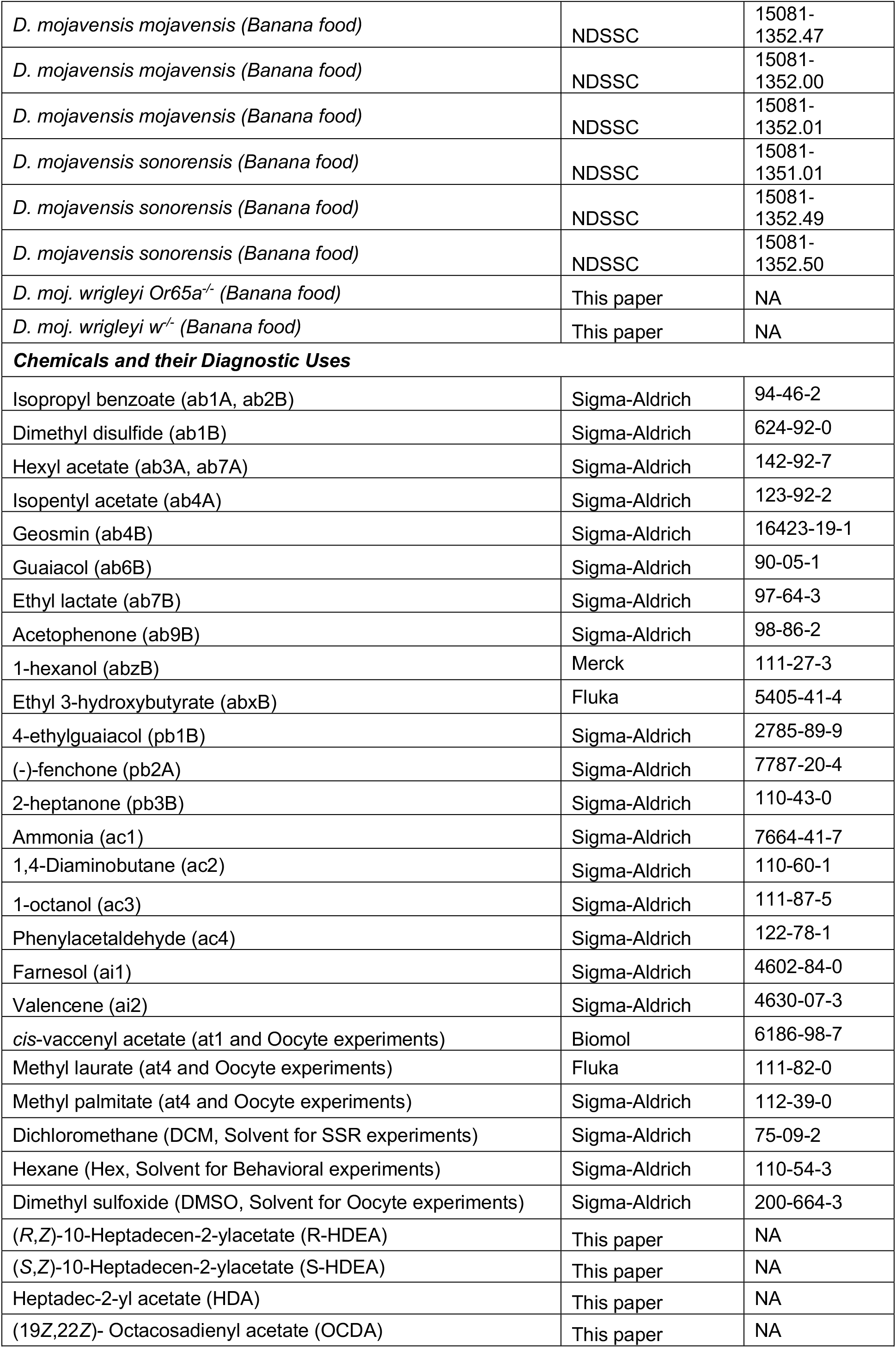
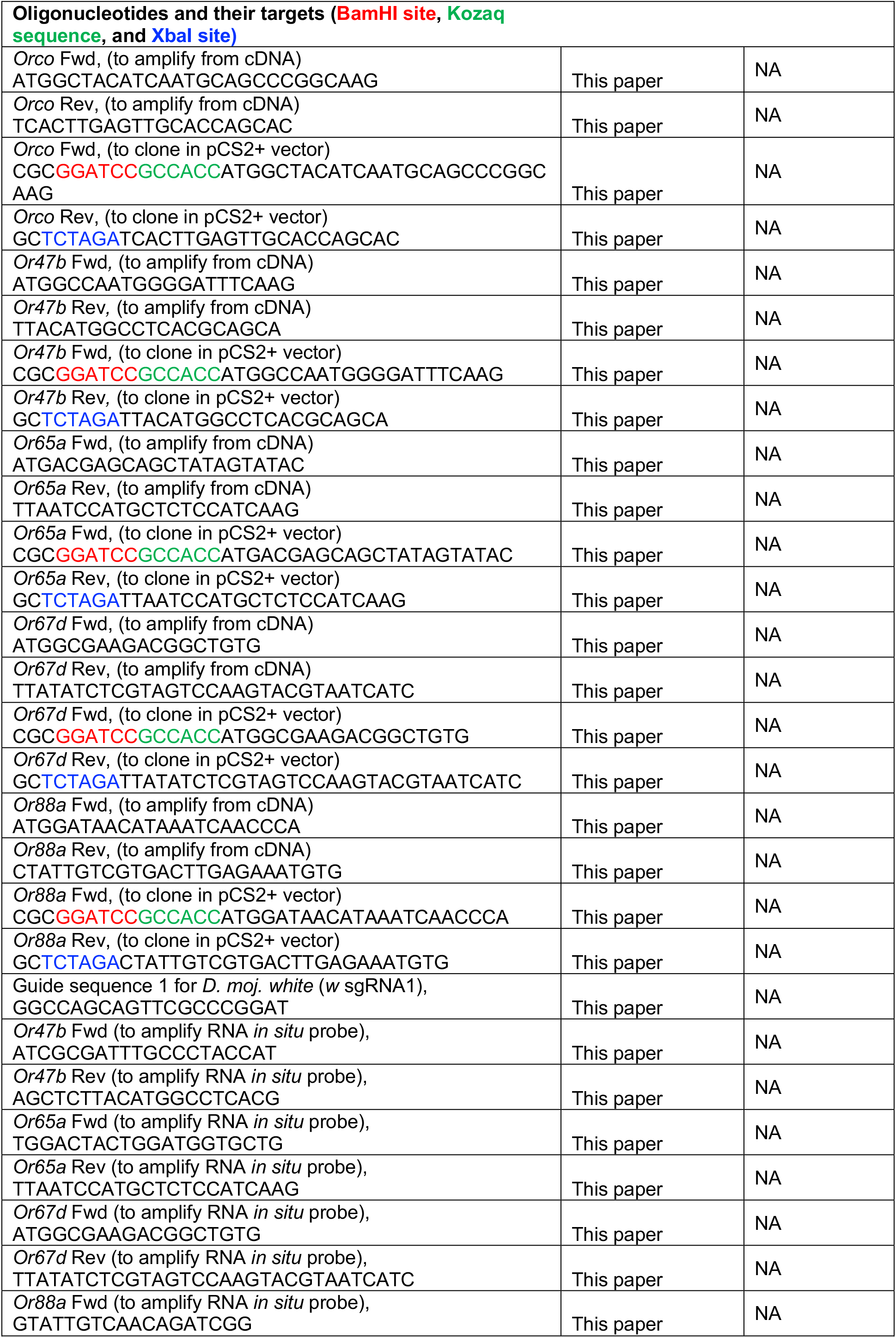
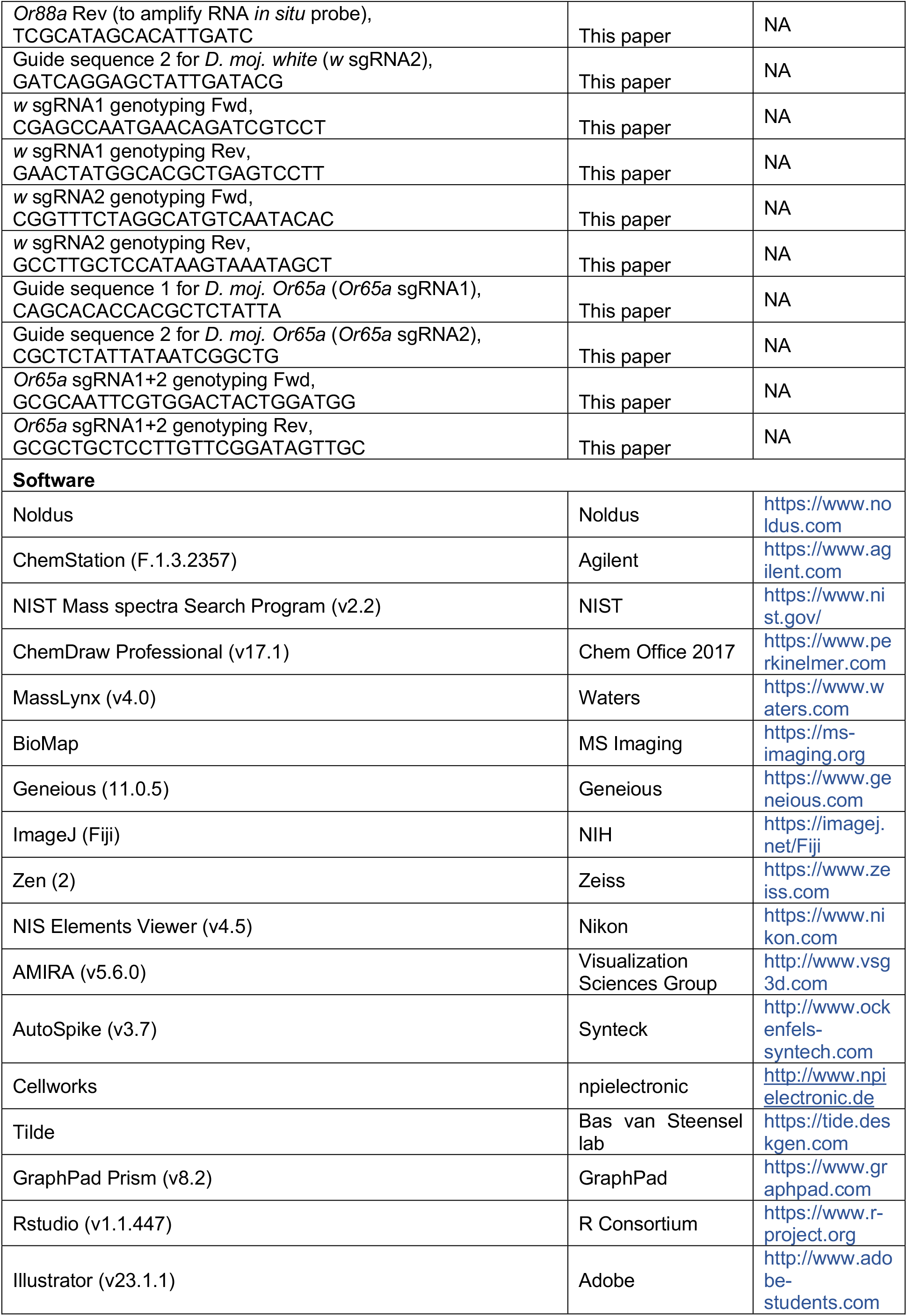
KEY RESOURCES TABLE.

**Figure Supplementary 1. Sexual behaviors of *D. mojavensis* subspecies.**

**A**, Schematic drawing of courtship behaviors of the four *D. mojavensis* subspecies. (**I**) orientation, (**II**) tapping with forelegs, (**III**) singing by wing fanning, (**IV**) licking the female genitals, (**V**) droplet discharge, and (**VI**) copulation attempt.

**B**, Courtship latency of males toward virgin consubspecific females in seconds (s) within a 20-minute time window. During the first minute, most males were eager to court by orienting and following the females. Among the four subspecies, *D. moj. wrigleyi* males exhibited shorter and less variable latencies to court. In this and the below panels, age of males and females is 10 days. Boxplots show the median, first and third quartile of the data. Different letters indicate significant differences between subspecies, Kruskal-Wallis test with Dunn’s post-hoc correction, n=52-74 per subspecies.

**C**, Percentage of males that released a fluidic droplet from their anus while courting the female. This panel reveals that more than half of the tested males while courting the female released a fluidic droplet from their anus. See Figure S2G for chemical analysis of these droplets. Courtship behaviors were recorded by GoPro Camera (Watch Movie S1-4 for details). Ns, Fisher’s exact test, n=10-20 per subspecies.

**D**, Courtship index [%] that males reveal toward their virgin consubspecific females. In response to the rapid male courtship elements, females slow down their movement, quiver their abdomen, vibrate their wings, and scissor them as signs for acceptance, while kicking with legs or accelerating the movement speed to signal rejection (Watch Movie S1-4). Overall, courtship rituals were comparable among the four subspecies. Ns *P* > 0.05, Kruskal-Wallis test with Dunn’s post-hoc correction, n=20.

**E**, Copulation latency in seconds (s). Males exhibiting no courtship behavior were excluded from analysis. *D. moj. wrigleyi* males exhibited shorter and less variable latencies to copulate. Kruskal-Wallis test with Dunn’s post-hoc correction, n=20-53 per subspecies.

**F**, Copulation success [%] of virgin couples in the different *D. mojavensis* subspecies within a 20-minute time window. Among the four subspecies, *D. moj. wrigleyi* males exhibited a higher percentage to be accepted for copulation. Fisher’s exact test, n=75.

**G**, Copulation duration of *D. mojavensis* subspecies in seconds (s). Unlike the prolonged copulation time in *D. melanogaster* (≥ 15 min) (Markow, 1996), copulation lasts for ~ two to three minutes in the *D. mojavensis* subspecies. Kruskal-Wallis test with Dunn’s post-hoc correction, n=20-53 per subspecies.

**Figure Supplementary 2. Chemical analysis of *D. mojavensis* subspecies.**

**A**, Representative gas chromatograms of hexane-body wash of 10-day-old male (virgin) and female flies (virgin and mated) (n=5). Colored peaks are the male-transferred compounds during mating; red, R&S–HDEA; blue, HDA; light green, OCDA.

**B**, Extracted ion chromatogram of hexane body wash of 10-day-old virgin male at the qualifier ion *m/z* 236 processed by chiral column (n=3). Colored peaks are S (brown) and R (red) enantiomers of 10*Z-* heptadecen-2yl acetate (R and S-HDEA; *m/z* 236).

**C**, Amount percentage of R and S-HDEA enantiomers in *D. moj. wrigleyi* and *D. moj. mojavensis*. Ns *P* > 0.01, Mann Whitney U test, n=3.

**D**, Representative imaging mass spectrometry for OCDA (see STAR Methods for details) by MALDI-TOF technique (Left: schematic drawing) of the abdominal surfaces of a 10-day-old male (virgin) and female fly (virgin and mated) (n=3).

**E**, Representative MALDI-TOF mass spectra of male (virgin) and female flies (virgin and mated) at the qualifier ion *m/z* 487 [M+K]+ of OCDA.

**F**, Representative gas chromatogram of ejaculatory bulbs of *D. moj. wrigleyi* obtained by solvent-free TD-GC-MS (Top: schematic drawing) of 10-day-old males (virgin) (n=3 replicates, each contains 5 ejaculatory bulbs). TD-GC-MS analyses reveal that all three acetates are present in high amounts in the ejaculatory bulb. Colored peaks indicate the male-specific compounds.

**G**, Gas chromatograms of male-released droplets during courtship in *D. moj. wrigleyi and D. moj. sonorensis* (n=3). Chemical analysis revealed the presence of R&S-HDEA and HDA in *D. moj. wrigleyi* droplets but not in droplets of *D. moj. sonorensis*, while OCDA was absent in the droplets of both subspecies. Due to the absence of OCDA signal, the x-axis was shortened. Colored peaks indicate the male-specific acetates (R&S-HDEA and HDA).

**Figure Supplementary 3. OCDA-induce courtship suppression, while the other acetates do not elicit attraction.**

**A**, Courtship index [%] of males towards dead consubspecific females perfumed with hexane as a control (black) or one of the male-specific acetates diluted in hexane (colored). See Figure 3B for details.

**B**, Headspace collection by SPME (see schematic drawing) and corresponding representative gas chromatogram from 10-day-old males of *D. moj. wrigleyi* trapped inside a mesh (dashed line) in a vial. Enlarged window represents R&S-HDEA (red) and HDA (blue) (n=3 replicates from each 20 males). Due to the absence of OCDA signal, the x-axis was shortened.

**C**, Representative tip recording traces from foreleg-tarsi using DMSO or OCDA (1 μg diluted in DMSO). Scale bar represents 150 milliseconds (ms).

**D**, Tip recording measurements from foreleg-tarsi of *D. moj. wrigleyi* and *D. moj. sonorensis* using R or S-HDEA, HDA, OCDA. Ns *P* > 0.05; ** *P* < 0.01, Kruskal-Wallis test followed by Dunn’s multiple comparisons test, n=5.

**E**, Schematics of wind tunnel paradigm (i.e., long-range attraction assay) where a group of five flies is released at the release platform and their landings are recorded.

**E’**, Boxplots of landing numbers per replicate toward R-HDEA (red), S-HDEA (brown), HDA (blue), OCDA (light green) or a mixture of the previous four compounds. None of the male acetates elicited any attraction, neither alone nor in a mixture of all compounds. Ns *P* > 0.05, Kruskal-Wallis test followed by Dunn’s multiple comparisons test, n=10.

**E’’**, Boxplots of landing numbers per replicate toward cactus juice in presence of hexane (black), R-HDEA (red), S-HDEA (brown), HDA (blue), OCDA (light green) or a mixture of the previous four compounds. Flies showed no increase in attraction between the cactus juice laced with the acetates singly or in mixture and the juice laced with hexane. Ns *P* > 0.05, Kruskal-Wallis test followed by Dunn’s multiple comparisons test, n=20.

**F,** Schematics of t-maze assay (i.e., short-range attraction assay) where a group of twenty-hours-starved flies is released at the starting point to choose between two arms.

**F’,** Boxplots of flies’ attraction index tested with cactus juice, R or S-HDEA, HDA, OCDA or a mixture of the previous four compounds vs water as a solvent. None of the acetates elicited any attraction, neither alone nor in a mixture. Ns *P* > 0.05; ** *P* < 0.01, Student t-test, n=10 replicates from each 30 flies.

**F**”, Boxplots of flies’ attraction index tested with cactus juice vs. the different male-specific compounds mixed with cactus juice. Flies showed no preference when cactus juice laced with the acetates singly or in mixture, compared to juice alone. Ns *P* > 0.05, Student t-test, n=10 replicates from each 30 flies.

**Figure Supplementary 4. Conserved detection mechanism of R-HDEA**

**A**, Copulation success [%] of *D. moj. wrigleyi* and *D. moj. sonorensis* males perfumed with hexane or OCDA diluted in hexane. See Figure 4A for details.

**B**, Copulation latency of the same males perfumed with hexane or OCDA diluted in hexane. See Figure 4B for details.

**C**, Competition between two consubspecific males, perfumed with OCDA or with hexane. See Figure 4C for details.

**D**, Top: competition between two males of *D. moj. mojavensis*, to mate with a virgin female. *\*\*\* P* < 0.001, chi-square test, n=25. Bottom: competition between two males of *D. moj. baja*, perfumed with R-HDEA or hexane, to copulate with a virgin consubspecific female. Ns *P* > 0.05, chi-square test, n=20.

**E**, Dose-response relationships for at4 neurons of *D. moj. wrigleyi* females (light grey) and males (dark grey) toward R-HDEA (Mean ± SEM). Two-way ANOVA followed by Sidek’s multiple comparison test between the two sexes responses to the same stimulus, ns P > 0.05; n=5-6 neurons.

**F**, Single-sensillum recording (SSR) measurements from all types of olfactory sensilla on antenna and maxillary palp, with OCDA (10 μg) as a stimulus. ab, antennal basiconic sensilla; ac, antennal coeloconic; at, antennal trichoid; ai, antennal intermediate; pb, palp basiconic (n=3-6).

**G**, Representative SSR traces from at4 sensillum of *D. moj. wrigleyi* and *D. moj. sonorensis* to DCM (as solvent), R-HDEA, S-HDEA, HDA, and OCDA (100 μg). Scale bar represents stimulus duration (0.5 second).

**H**, Responses of at4 sensilla in *D. mojavensis* subspecies (color coded as in Figure 1A) to R-HDEA (1000 μg). Ns *P* > 0.01, Mann Whitney U test, n=6.

**I**, Responses of the *D. melanogaster* at1 (black) and at4 (grey) to R, HDEA, S-HDEA, HDA, OCDA, methyl palmitate (MP; diagnostic odor for Or88a), methyl laurate (ML; diagnostic odor for Or47b) and *cis* vaccenyl acetate (*c*VA; diagnostic odor for Or67d) (10 μg diluted in DCM). SSR amylases reveal that none of the *D. moj. wrigleyi* male-specific acetates elicited any response in the at1 nor at4 sensillum of *D. melanogaster*. Filled circles in this panel indicate significant difference from solvent responses. Ns *P* > 0.05; ** *P* < 0.01; *** *P* < 0.001, Mann Whitney U test, n=3-4.

**J**, Alignments sequences of *D. moj. wrigleyi-OR65a* and *D. melanogaster-OR65a/b/c* amino acid sequences. Blue letters represent the similarities while red letters represent the polymorphic sites.

**Figure Supplementary 5. Functional characterization of sex pheromone receptors and generation of white mutant flies.**

**A**, Phylogenetic analysis of pheromone receptors (OR67d, OR88a, Or65 and OR47b) in *D. melanogaster* and *D. moj. wrigleyi* using Or65a ortholog in *Glossina morsitans* as an outgroup. Of the six pheromone sensing receptors in *D. melanogaster*, five are present in the *D. moj. wrigleyi* genome (Guo and Kim, 2007): GI19867 and GI19869 (OR47b1 and OR47b2, respectively), GI12096 (OR65a/b/c), GI11463 (OR67d), and GI23341 (OR88a). GI12096 has three paralogs in *D. melanogaster* (OR65a/b/c) and shares the highest degree of protein alignment with *D. melnaogaster-OR65a* (Guo and Kim, 2007) compared to others (Figure S4J). The scale bar for branch length represents the number of substitutions per site.

**B**, Responses of the five odorant receptors (indicated in different colors), heterologously expressed in *X. laevis* oocytes to S-HDEA, HDA, methyl laurate (diagnostic odor for Or47b), and methyl palmitate (diagnostic odor for Or88a) (1mM diluted in DMSO). Filled circles in this and other panels indicate significant difference from solvent responses, ns *P* > 0.05; * *P* < 0.05; *** *P* < 0.001, Mann Whitney U test, n=2-8.

**C**, Responses of the *D. moj. wrigleyi/Or65a* gene heterologously expressed in *D. melanogaster* at1 to previous-mentioned compounds and cis-vaccenyl acetate (diagnostic odor for Or67d) (10 μg diluted in DCM). Ns *P* > 0.05; *** *P* < 0.001, Mann Whitney U test, n=3-4.

**D**, Responses of *Or65a* heterozygous (black) and homozygous (grey) animals to R-HDEA and methyl laurate (ML) (100 μg diluted in DCM). Filled-circles indicate significant difference between both groups. Ns *P* > 0.05; *** *P* < 0.001, Mann Whitney U test, n=6.

**E**, Schematic drawing of the structure of the white gene (*G/10968-6586041)* illustrating the strategy for generating knockouts using CRISPR-Cas9. The two guide RNAs sequences are shown below; scissors denote the cutting positions. All knockouts were validated by sequencing the targeted locus in homozygous mutants prior to establishing lines. The *white* gene knockout animals carry a 15 bp deletion that results in early termination.

**F**, A lateral macrograph of wildtype and *white* mutant female (right) or wildtype and *white* mutant male (left) of *D. moj. wrigleyi*. In addition to the eye color change, the yellowish color of the male’s accessory glands disappeared. Scale bar represents 500 μm.

**G**, Protein alignment of OR65a in the four *D. moj*. subspecies in an open reading frame. Blue letters represent the similarities while red letters represent the polymorphic sites.

**Figure Supplementary 6. Characterization of antennal lobe glomeruli in *D. mojavensis* subspecies.**

**A**, Normalized volumes of 54 glomeruli (out of 57) for *D. moj. wrigleyi* and *D. moj. sonorensis* females. Glomeruli in red are *D. mojavensis-specific* novel glomeruli, which named according to their relative position to the adjacent glomeruli. Filled bars indicate significant differences between both subspecies. Two-way ANOVA followed by Sidek’s multiple comparison test between the two subspecies’ responses to the same stimulus, ns P > 0.05; *** *P* < 0.001, n=3-4 animals per subspecies. See Table S1 and Figure S6A for more details.

**B**, A pattern of neurobiotin backfilled neurons (magenta) from at4 sensillum that reveals similar innervation in *D. moj. wrigleyi* (right) and *D. moj. sonorensis* (left) to VA8, VA1v and VA1d glomeruli.

**C**, Three-dimensional reconstruction of antennal lobes from representative female brains of *D. moj. wrigleyi, D. moj. sonorensis* and *D. melanogaster*. DA1, red; DL3, cyan; VL2a, black. Scale bar represents 20 μm.

**D**, Fluorescent staining for neurobiotin (green) and nc82 (magenta) in *D. moj. wrigleyi* antennal lobe backfilled from at1 sensillum (identified by electrophysiological recordings; Figure 4E). Backfilling of the at1 sensillum of *D. moj. wrigleyi* revealed a similar innervation target as in *D. melanogaster* (Couto et al., 2005) to the DA1 glomerulus but with two neuronal tracts innervating separately an anterior and posterior region of this glomerulus. The backfill image corresponds to a projection of 28 Z-stacks (Watch Movie S9).

**E**, Reconstructions for the backfill signal that innervate the DA1 (red) but not DL3 (cyan) glomerulus.

**F**, Representative SSR traces from at1 sensillum of *D. moj. wrigleyi* DCM, *cis*-vaccenyl acetate (cVA, 100 μg). Consistent with the innervation pattern of *D. mojavensis* at1 neurons, SSR analysis of the at1 sensillum revealed the presence of at least two OSNs in this sensillum type (Prieto-Godino et al., 2019). Scale bar represents the stimulus duration (0.5 second).

**G**, Antennal lobe volumes (μm^3^) for males (black) and females (grey) of *D. moj. wrigleyi* and *D. moj. sonorensis*. Filled circles in this and below panels indicate significant difference from the other sex within the same species, ns *P* > 0.05; * *P* < 0.05; ** *P* < 0.01, Mann Whitney U test, n=4-6.

**H**, Normalized volumes of DA1, VL2a and DL3 for *D. moj. wrigleyi* and *D. moj. sonorensis* females and males.

**Figure Supplementary 7. S-HDEA and HDA are not involved in sexual isolation among *D. mo%avensis* subspecies.**

**A**, Competition between two males of different subspecies, *D. moj. sonorensis* male perfumed with S-HDEA (red droplet, left panel) or HDA (blue droplet, right panel) and *D. moj. wrigleyi* male perfumed with hexane (black droplets), to copulate with *D. moj. wrigleyi* virgin female. Pie-charts in A-B represent copulation success [%] of the rival males, ** *P* < 0.01; *** *P* < 0.001, chi-square test, number of the replicates for is 21 and 20, respectively. All males and females used in this and other panels were 10-day-old virgin flies.

**B**, Competition between two males of different subspecies, *D. moj. sonorensis* male perfumed with S-HDEA (red droplet, left panel) or HDA (blue droplet, right panel) and *D. moj. wrigleyi* male perfumed with hexane (black droplets), to copulate with *D. moj. sonorensis* virgin female. Number of the replicates for is 21 for both panels.

**Table Supplementary 1.**
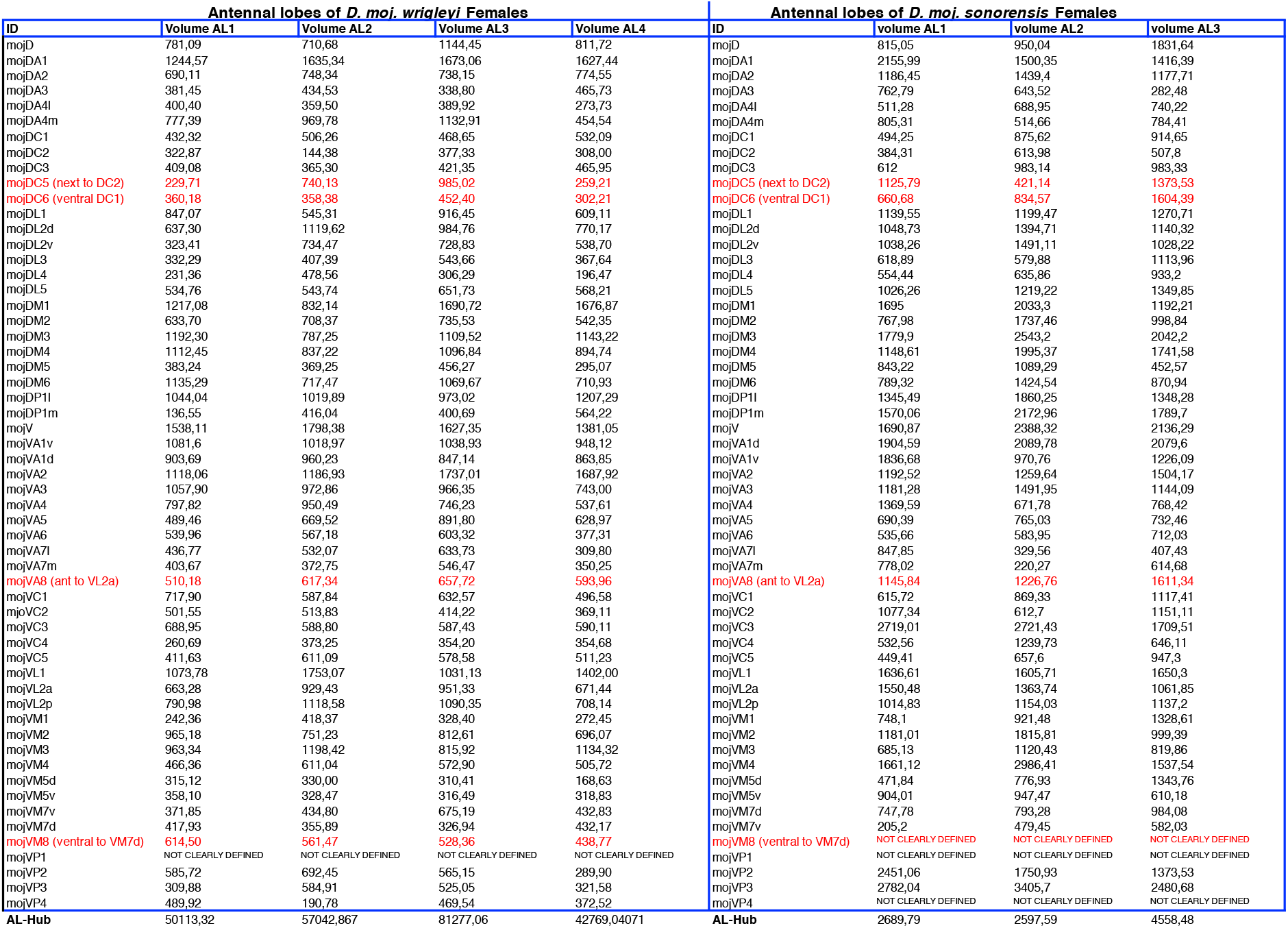
Comparison of antennal lobes in *D. moj. wrigleyi* and *D. moj. sonorensis*. 57 glomeruli were identified in both *D. mojavensis* subspecies. Identification of glomeruli was verified by comparing the reconstructed images to the map of *D. melanogaster* AL (Grabe et al., 2015). *D. mo/avensis*-specific glomeruli are written in red (i.e., they are not present in *D. melanogaster)*. The volumes of VP1 in both subspecies and VM8 and VP4 in *D. moj. sonorensis* could not be quantified.

**Movie Supplementary 1. Sexual behaviors of *D. mojavensis wrigleyi*.**

During the first minute, most males of the four subspecies were eager to court by orienting and following the females. Subsequently, males tapped the females’ bodies with their forelegs, followed by wing spreading and fanning for vibrational song production. Females responded to the males’ song by vibrating their wings. Males followed the females by extending their proboscis to lick the females’ genitalia and then attempted copulation. Males while courting the female in addition released a fluidic droplet from their anus. We called this novel trait “dropping behavior”. In response to the rapid male courtship elements, females slow down their movement, quiver their abdomen, vibrate their wings and scissor them as signs for acceptance, while kicking with legs or accelerating the movement speed to signal rejection.

**Movie Supplementary 2. Sexual behaviors of *D. moj. mojavensis*.**

**Movie Supplementary 3. Sexual behaviors of *D. moj. sonorensis*.**

**Movie Supplementary 4. Sexual behaviors of *D. moj. baja*.**

**Movie Supplementary 5. 3-D reconstruction of *D. moj. wrigleyi* antennal lobe (female).**

**Movie Supplementary 6. 3-D reconstruction of *D. moj. sonorensis* antennal lobe (female).**

**Movie Supplementary 7. Neurobiotin backfilled neurons from at4 sensillum in *D. moj. wrigleyi*.** Fluorescent staining for neurobiotin (green) and nc82 (magenta) of *D. moj. wrigleyi* antennal lobe, backfilled from at4 sensillum.

**Movie Supplementary 8. Neurobiotin backfilled neurons from at4 sensillum in *D. moj. sonorensis*.**

Fluorescent staining for neurobiotin (green) and nc82 (magenta) of *D. moj. sonorensis* antennal lobe, backfilled from at4 sensillum.

**Movie Supplementary 9. Neurobiotin backfilled neurons from at1 sensillum in *D. moj. wrigleyi*.** Fluorescent staining for neurobiotin (green) and nc82 (magenta) of *D. moj. wrigleyi* antennal lobe, backfilled from at1 sensillum.

**File Supplementary 1**. Alignments sequences of *Or47bl* and *Or47b2* loci in *D. moj. wrigleyi* (provided as Fasta file).

**File Supplementary 2.** 3-D Pdf of antennal lobe reconstruction of *D. moj. wrigleyi* (female).

**File Supplementary 3.** 3-D Pdf of antennal lobe reconstruction of *D. moj. sonorensis* (female).

## References

Ahmed, O.M., Avila-Herrera, A., Tun, K.M., Serpa, P.H., Peng, J., Parthasarathy, S., Knapp, J.M., Stern, D.L., Davis, G.W., Pollard, K.S., et al. (2019). Evolution of mechanisms that control mating in *Drosophila* males. Cell Rep 27, 2527–2536 e2524.

Allan, C.W., and Matzkin, L.M. (2019). Genomic analysis of the four ecologically distinct cactus host populations of *Drosophila mojavensis*. Bmc Genomics 20, 732.

Arguello, J.R., and Benton, R. (2017). Open questions: Tackling Darwin’s “instincts”: the genetic basis of behavioral evolution. Bmc Biol 15.

Auer, T.O., and Benton, R. (2016). Sexual circuitry in *Drosophila*. Curr Opin Neurobiol 38, 18–26.

Becher, P.G., Bengtsson, M., Hansson, B.S., and Witzgall, P. (2010). Flying the fly: long-range flight behavior of *Drosophila melanogaster* to attractive odors. J Chem Ecol 36, 599–607.

Benton, R., Vannice, K.S., Gomez-Diaz, C., and Vosshall, L.B. (2009). Variant ionotropic glutamate receptors as chemosensory receptors in *Drosophila*. Cell 136, 149–162.

Bischof, J., Maeda, R.K., Hediger, M., Karch, F., and Basler, K. (2007). An optimized transgenesis system for *Drosophila* using germ-line-specific phi C31 integrases. P Natl Acad Sci USA 104, 3312–3317.

Brinkman, E.K., Chen, T., Amendola, M., and van Steensel, B. (2014). Easy quantitative assessment of genome editing by sequence trace decomposition. Nucleic Acids Res 42, e168.

Chin, J.S.R., Ellis, S.R., Pham, H.T., Blanksby, S.J., Mori, K., Koh, Q.L., Etges, W.J., and Yew, J.Y. (2014). Sex-specific triacylglycerides are widely conserved in *Drosophila* and mediate mating behavior. Elife 3.

Couto, A., Alenius, M., and Dickson, B.J. (2005). Molecular, anatomical, and functional organization of the *Drosophila* olfactory system. Curr Biol 15, 1535–1547.

Crowley-Gall, A., Date, P., Han, C., Rhodes, N., Andolfatto, P., Layne, J.E., and Rollmann, S.M. (2016). Population differences in olfaction accompany host shift in *Drosophila mojavensis*. P Roy Soc B-Biol Sci 283.

Cvacka, J., and Svatos, A. (2003). Matrix-assisted laser desorption/ionization analysis of lipids and high molecular weight hydrocarbons with lithium 2,5-dihydroxybenzoate matrix. Rapid Commun Mass Sp 17, 2203–2207.

Date, P., Dweck, H.K.M., Stensmyr, M.C., Shann, J., Hansson, B.S., and Rollmann, S.M. (2013). Divergence in olfactory host plant preference in *D& mojavensis* in response to cactus host use. Plos One 8.

Dekker, T., Ibba, I., Siju, K.P., Stensmyr, M.C., and Hansson, B.S. (2006). Olfactory shifts parallel superspecialism for toxic fruit in *Drosophila melano9as#er* sibling, *D. sechellia*. Curr Biol 16, 101–109.

Ding, Y., Berrocal, A., Morita, T., Longden, K.D., and Stern, D.L. (2016). Natural courtship song variation caused by an intronic retroelement in an ion channel gene. Nature 536, 329–+.

Dobzhansky, T.H. (1937). Genetics and the origin of species (New York, USA: Columbia University Press).

Dweck, H.K.M., Ebrahim, S.A.M., Khallaf, M.A., Koenig, C., Farhan, A., Stieber, R., Weissflog, J., Svatos, A., Grosse-Wilde, E., Knaden, M., et al. (2016). Olfactory channels associated with the *Drosophila* maxillary palp mediate short- and long-range attraction. Elife 5.

Dweck, H.K.M., Ebrahim, S.A.M., Thoma, M., Mohamed, A.A.M., Keesey, I.W., Trona, F., Lavista-Llanos, S., Svatos, A., Sachse, S., Knaden, M., et al. (2015). Pheromones mediating copulation and attraction in *Drosophila*. P Natl Acad Sci USA 112, E2829–E2835.

Edgar, R.C. (2004). MUSCLE: multiple sequence alignment with high accuracy and high throughput. Nucleic Acids Res 32, 1792–1797.

Ejima, A., Smith, B.P.C., Lucas, C., Van Naters, W.V., Miller, C.J., Carlson, J.R., Levine, J.D., and Griffith, L.C. (2007). Generalization of courtship learning in *Drosophila* is mediated by cis-vaccenyl acetate. Curr Biol 17, 599–605.

Endler, J.A. (1992). Signals, Signal Conditions, and the Direction of Evolution. Am Nat 139, S125–S153.

Etges, W.J. (2019). Evolutionary genomics of host plant adaptation: insights from *Drosophila*. Curr Opin Insect Sci 36, 96–102.

Etges, W.J., and Ahrens, M.A. (2001). Premating isolation is determined by larval-rearing substrates in cactophilic *Drosophila mojavensis.* V. Deep geographic variation in epicuticular hydrocarbons among isolated populations. Am Nat 158, 585–598.

Etges, W.J., de Oliveira, C.C., Gragg, E., Ortiz-Barrientos, D., Noor, M.A.F., and Ritchie, M.G. (2007). Genetics of incipient speciation in *Drosophila mojavensis.* I. Male courtship song, mating success, and genotype x environment interactions. Evolution 61, 1106–1119.

Etges, W.J., Over, K.F., De Oliveira, C.C., and Ritchie, M.G. (2006). Inheritance of courtship song variation among geographically isolated populations of *Drosophila mojavensis*. Anim Behav 71, 1205–1214.

Fishilevich, E., and Vosshall, L.B. (2005). Genetic and functional subdivision of the *Drosophila* antennal lobe. Curr Biol 15, 1548–1553.

Gonzalez, F., Witzgall, P., and Walker, W.B. (2016). Protocol for heterologous expression of insect odourant receptors in *Drosophila*. Frontiers in Ecology and Evolution 4.

Grabe, V., Baschwitz, A., Dweck, H.K.M., Lavista-Llanos, S., Hansson, B.S., and Sachse, S. (2016). Elucidating the neuronal architecture of olfactory glomeruli in the *Drosophila* antennal lobe. Cell Rep 16, 3401–3413.

Grabe, V., Strutz, A., Baschwitz, A., Hansson, B.S., and Sachse, S. (2015). Digital In vivo 3D atlas of the antennal lobe of *Drosophila melano9as#er*. J Comp Neurol 523, 530–544.

Guindon, S., Dufayard, J.F., Lefort, V., Anisimova, M., Hordijk, W., and Gascuel, O. (2010). New algorithms and methods to estimate maximum-likelihood phylogenies: assessing the performance of PhyML 3.0. Syst Biol 59, 307–321.

Guo, S., and Kim, J. (2007). Molecular evolution of *Drosophila* odorant receptor genes. Mol Biol Evol 24, 1198–1207.

Hansson, B.S., and Stensmyr, M.C. (2011). Evolution of insect olfaction. Neuron 72, 698–711.

Heed, W.B. (1978). Ecology and genetics of sonoran desert *Drosophila* (Proceedings in Life Sciences. Springer, New York, NY).

Horka, P., Vrkoslav, V., Hanus, R., Peckova, K., and Cvacka, J. (2014). New MALDI matrices based on lithium salts for the analysis of hydrocarbons and wax esters. J Mass Spectrom 49, 628–638.

Kaftan, F., Vrkoslav, V., Kynast, P., Kulkarni, P., Bocker, S., Cvacka, J., Knaden, M., and Svatos, A. (2014). Mass spectrometry imaging of surface lipids on intact *Drosophila melanogaster* flies. J Mass Spectrom 49, 223–232.

Katoh, K., and Standley, D.M. (2013). MAFFT multiple sequence alignment software version 7: Improvements in performance and usability. Mol Biol Evol 30, 772–780.

Knowles, L.L., and Markow, T.A. (2001). Sexually antagonistic coevolution of a postmating-prezygotic reproductive character in desert *Drosophila*. Proc Natl Acad Sci USA 98, 8692–8696.

Kohl, J., Huoviala, P., and Jefferis, G.S.X.E. (2015). Pheromone processing in *Drosophila*. Curr Opin Neurobiol 34, 149–157.

Kohl, J., Ostrovsky, A.D., Frechter, S., and Jefferis, G.S.X.E. (2013). A bidirectional circuit switch reroutes pheromone signals in male and female brains. Cell 155, 1610–1623.

Kondoh, Y., Kaneshiro, K.Y., Kimura, K., and Yamamoto, D. (2003). Evolution of sexual dimorphism in the olfactory brain of Hawaiian *Drosophila*. P Roy Soc B-Biol Sci 270, 1005–1013.

Krebs, R.A., and Markow, T.A. (1989). Courtship behavior and control of reproductive isolation in *Drosophila mojavensis*. Evolution 43, 908–913.

Kurtovic, A., Widmer, A., and Dickson, B.J. (2007). A single class of olfactory neurons mediates behavioural responses to a *Drosophila* sex pheromone. Nature 446, 542–546.

Lande, R. (1981). Models of speciation by sexual selection on polygenic traits. P Natl Acad Sci USA 78, 3721–3725.

LaRue, K.M., Clemens, J., Berman, G.J., and Murthy, M. (2015). Acoustic duetting in *Drosophila virilis* relies on the integration of auditory and tactile signals. Elife 4.

Lebreton, S., Grabe, V., Omondi, A.B., Ignell, R., Becher, P.G., Hansson, B.S., Sachse, S., and Witzgall, P. (2014). Love makes smell blind: mating suppresses pheromone attraction in *Drosophila* females via Or65a olfactory neurons. Sci Rep-Uk 4.

Linz, J., Baschwitz, A., Strutz, A., Dweck, H.K.M., Sachse, S., Hansson, B.S., and Stensmyr, M.C. (2013). Host plant-driven sensory specialization in *Drosophila erecta*. P Roy Soc B-Biol Sci 280.

Liu, W.W., Liang, X.H., Gong, J.X., Yang, Z., Zhang, Y.H., Zhang, J.X., and Rao, Y. (2011). Social regulation of aggression by pheromonal activation of Or65a olfactory neurons in *Drosophila*. Nat Neurosci 14, 896–U119.

Manoli, D.S., Foss, M., Villella, A., Taylor, B.J., Hall, J.C., and Baker, B.S. (2005). Male-specific fruitless specifies the neural substrates of *Drosophila* courtship behaviour. Nature 436, 395–400.

Markow, T.A. (1991). Sexual isolation among populations of *Drosophila mojavensis*. Evolution 45, 1525–1529.

Markow, T.A. (1996). Evolution of *Drosophila* mating systems. Evol Biol 29, 73–106.

Markow, T.A., and O’Grady, P.M. (2005). Evolutionary genetics of reproductive behavior in *Drosophila:* connecting the dots. Annual Review of Genetics 39, 263–291.

Matzkin, L.M. (2014). Ecological genomics of host shifts in *Drosophila mojavensis*. Ecological genomics: ecology and the evolution of genes and genomes 781, 233–247.

Matzkin, L.M., and Markow, T.A. (2013). Transcriptional differentiation across the four cactus host races of Drosophila mojavensis. (New York, NY: Nova Science Publishers).

Mayr, E. (1942). Systematics and the origin of species (Columbia University Press).

Mercier, D., Tsuchimoto, Y., Ohta, K., and Kazama, H. (2018). Olfactory landmark-based communication in interacting *Drosophila*. Curr Biol 28, 2624–+.

Moon, S.J., Köttgen, M., Jiao, Y., Xu, H., and Montell, C. (2006). A taste receptor required for the caffeine response in vivo. Curr Biol 16, 1812–1817.

Nemeth, D.C., Ammagarahalli, B., Layne, J.E., and Rollmann, S.M. (2018). Evolution of coeloconic sensilla in the peripheral olfactory system of *Drosophila mojavensis*. J Insect Physiol 110, 13–22.

Newby, B.D., and Etges, W.J. (1998). Host preference among populations of *Drosophila mojavensis* (Diptera: Drosophilidae) that use different host cacti. J Insect Behav 11, 691–712.

Nosil, P. (2012). Ecological speciation. Ox Ecol Ev, 1–280.

O’Grady, P.M., and Markow, T.A. (2012). Rapid morphological, behavioral, and ecological evolution in *Drosophila:* comparisons between the endemic Hawaiian *Drosophila* and the cactophilic repleta species group. Rapidly evolving genes and genetic systems 1, 176–186.

Olsson, S.B., and Hansson, B.S. (2013). Electroantennogram and single sensillum recording in insect antennae. Methods Mol Biol 1068, 157–177.

Pfeiler, E., Castrezana, S., Reed, L.K., and Markow, T.A. (2009). Genetic, ecological and morphological differences among populations of the cactophilic *Drosophila mojavensis* from southwestern USA and northwestern Mexico, with descriptions of two new subspecies. J Nat Hist 43, 923–938.

Prieto-Godino, L.L., Rytz, R., Cruchet, S., Bargeton, B., Abuin, L., Silbering, A.F., Ruta, V., Dal Peraro, M., and Benton, R. (2017). Evolution of acid-sensing olfactory circuits in drosophilids. Neuron 93, 661–+.

Prieto-Godino, L.L., Silbering, A.F., Khallaf, M.A., Cruchet, S., Bojkowska, K., Pradervand, S., Hansson, B.S., Knaden, M., and Benton, R. (2019). Functional integration of “undead” neurons in the olfactory system. bioRxiv. http://dx.doi.org/10.1101/623488.

Ritchie, M.G. (2007). Sexual selection and speciation. Annual Review of Ecology, Evolution, and Systematics 38, 79–102.

Ruiz, A., Heed, W.B., and Wasserman, M. (1990). Evolution of the *mojavensis* cluster of cactophilic *Drosophila* with descriptions of 2 new species. J Hered 81, 30–42.

Ruta, V., Datta, S.R., Vasconcelos, M.L., Freeland, J., Looger, L.L., and Axel, R. (2010). A dimorphic pheromone circuit in *Drosophila* from sensory input to descending output. Nature 468, 686–U106.

Saina, M., and Benton, R. (2013). Visualizing olfactory receptor expression and localization in *Drosophila*. Methods Mol Biol 1003, 211–228.

Seehausen, O., Terai, Y., Magalhaes, I.S., Carleton, K.L., Mrosso, H.D., Miyagi, R., van der Sluijs, I., Schneider, M.V., Maan, M.E., Tachida, H., et al. (2008). Speciation through sensory drive in cichlid fish. Nature 455, 620–626.

Seeholzer, L.F., Seppo, M., Stern, D.L., and Ruta, V. (2018). Evolution of a central neural circuit underlies *Drosophila* mate preferences. Nature 559, 564–+.

Smadja, C., and Butlin, R.K. (2009). On the scent of speciation: the chemosensory system and its role in premating isolation. Heredity 102, 77–97.

Spieth, H.T. (1952). Mating behavior within the genus *Drosophila* (Diptera). B Am Mus Nat Hist 99, 401–474.

Spieth, H.T. (1974). Courtship Behavior in *Drosophila*. Annu Rev Entomol 19, 385–405.

Steele, R.H. (1986). Courtship feeding in *Drosophila subobs*.1. The Nutritional significance of courtship feeding. Anim Behav 34, 1087–1098.

Stokl, J., Strutz, A., Dafni, A., Svatos, A., Doubsky, J., Knaden, M., Sachse, S., Hansson, B.S., and Stensmyr, M.C. (2010). A deceptive pollination system targeting drosophilids through olfactory mimicry of yeast. Curr Biol 20, 1846–1852.

Symonds, M.R.E., and Wertheim, B. (2005). The mode of evolution of aggregation pheromones in *Drosophila* species. J Evolution Biol 18, 1253–1263.

van Naters, W.V.G., and Carlson, J.R. (2007). Receptors and neurons for fly odors in *Drosophila*. Current Biology 17, 606–612.

Waterhouse, R.M., Seppey, M., Simao, F.A., Manni, M., Ioannidis, P., Klioutchnikov, G., Kriventseva, E.V., and Zdobnov, E.M. (2018). BUSCO applications from quality assessments to gene prediction and phylogenomics. Mol Biol Evol 35, 543–548.

Yew, J.Y., Dreisewerd, K., Luftmann, H., Muthing, J., Pohlentz, G., and Kravitz, E.A. (2009). A new male sex pheromone and novel cuticular cues for chemical communication in *Drosophila*. Current Biology 19, 1245–1254.

Zawistowski, S., and Richmond, R.C. (1986). Inhibition of courtship and mating of *Drosophila melanogaster* by the male-produced lipid, cis vaccenyl acetate. J Insect Physiol 32, 189–192.

Zhang, D.D., and Lofstedt, C. (2013). Functional evolution of a multigene family: orthologous and paralogous pheromone receptor genes in the turnip moth, *Agrotis segetum*. Plos One 8.

Zouros, E., and Dentremont, C.J. (1980). Sexual isolation among populations of *Drosophila mojavensis* - response to pressure from a related species. Evolution 34, 421–430.

