## Supplementary text for "Mate discrimination among subspecies through a conserved olfactory pathway" (from another version of this preprint: version 1, 2019-11-25)

**Chemical identification and synthesis**

**Chemical identification**

TDU-GC/EIMS data obtained from virgin males and females show three chromatographic peaks abundant in males. The first one (KI 1998) was the most abundant. Molecular ion peak is not visible, but rather *m/z* 236 is abundant. Accurate mass provided molecular composition C_17_H_32_. A large EE ion series of 194 (236-C3H6), 180, 166 up to 82 (consecutive losses of CH_2_) indicate unsaturation in alicyclic chain. Diagnostic peaks at *m/z* 87 (α-cleavage, C_4_H_7_O_2_), 61 (C_2_H_3_O_2_) 43 indicate acetyl group on position 2. Then *m/z* 236 peak represents [M-CH_3_COOH]^+●^. The double bond position was determined by converting the compound to dimethyldisulfite (DMDS) adduct^1^. The EIMS spectrum show *m/z* 390 [M]^+●^ and indicative fragments A: *m/z* 245 and *m/z* 185 [A-CH_3_COOH]^+^; B: 145 plus additional ions at *m/z* 87 and 61. Data analysis suggests 10-heptadecen-2-yl-acetate. The prepared (*rac,*10*Z*)-heptadecen-2-yl-acetate (HDEA) show undistinguishable EIMS spectra and chromatographic retention as the natural product. Chirality and enantiomeric purity of natural acetate was determined on a chiral GC column using prepared (*2R,*10*Z*)-heptadecen-2-yl-acetate and (2R*,*10*Z*)-heptadecen-2-yl-acetate standards.

The second peak with longer retention (KI 2017) show in EIMS spectrum low-abundant molecular ion *m/z* 298 M^+●^, *m/z* 256 [M-C_2_H_2_O]^+●^ and intense *m/z* 238 [M- CH_3_COOH]^+●^. In addition, OE ion series from *m/z* 153-55 separated by 14Da and peaks at *m/z* 87 (α-cleavage, C_4_H_7_O_2_), 61 (C_2_H_3_O_2_) 43 indicating saturated aliphatic chain and acetyl group on position 2. The second peak is heptadec-2-yl-acetate (HDA). HDA have also chiral center at position 2, however 2*R*- and 2*S*-enantiomers were present in 50/50 ratio and only and *rac-*heptadec-2-yl-acetate was prepared and further used in this study.

Third peak with much longer retention (KI 3058) shows in abundant molecular ion *m/z* 448 (M^+●^, accurate mass *m/z* 448.42951 for C_30_H_56_O_2_ calculated 448.428032) and fragment ions at *m/z* 405 [M-C_2_H_2_O]^+●^ and *m/z* 388 [M- CH_3_COOH]^+●^ accompanied with EE ion series from *m/z* 264-166 OE series *m/z* 149-67 both spaced by 14Da. Acetate function was deduced from *m/z* 61 (C_2_H_3_O_2_) most probably in position 1. Intense ions *m/z* 149, 135, 121 and 109 indicate omega-6 and omega-9 skipped diene structure. Data suggest (*Z,Z*)-19,22-octacosadien-1-yl acetate (OCDA) as the third peak and the proposed structure was confirmed by chemical synthesis.

**Chemical synthesis**

Commercially available chemicals were used without further purification. Tetrahydrofuran (anhydrous, Sigma-Aldrich) and dichloromethane (Roth) were used as received. All reactions except the enzymatic hydrolysis were carried out under an atmosphere of argon. Preparative column chromatography was performed on Silica gel 60 (230-400 mesh, Carl Roth GmbH) and TLC analysis on commercial Merck silica gel 60 F_254_ plates. NMR spectra were measured on a Bruker Avance 400 NMR spectrometer. Chemical shifts are reported in ppm downfield from TMS.

**Synthesis of (*R*) and (*S*,*Z*)-10-Heptadecen-2-ylacetate (R&S-HDEA).**

*Synthesis of (Z)-10-Heptadecen-2-one.*

Palmitoleic acid (250 mg, 0.98 mmol) was dissolved in 10 ml of THF and cooled to -78°C. A 1.6 M solution of MeLi in diethylether (1.53 ml, 2.45 mmol) was added dropwise and the mixture was allowed to warm to room temperature by removing the cooling bath. The reaction was monitored via TLC. Upon completion (after ca. 2.5h) the mixture was poured into 20 ml of 1N HCl at 0°C and extracted with diethylether (3x15 ml). The combined organic layers were washed with saturated NaHCO_3_ (20 ml) and brine (20 ml), dried with anhydrous NaSO_4_, filtered and concentrated in vacuo. After purification via silica gel column chromatography (12:1 hexane/EtOAc) (*Z*)-10-Heptadecen-2-one was obtained as colourless oil (225 mg, 91%).^1^H-NMR: (400 MHz, CDCl_3_) δ = 5.35 (m, 2H), 2.41 (t, *J* = 7.3 Hz, 2H), 2.13 (s, 3H), 2.01 (q, *J* = 5.7 Hz, 4H), 1.57 (m, 2H), 1.29 (m, 16H), 0.89 (t, *J* = 6.0 Hz, 3H)ppm; ^13^C-NMR: (100 MHz, CDCl_3_) δ = 209.2, 130.0, 129.7, 43.8, 31.8, 29.8, 29.72, 29.68, 29.28, 29.15, 29.09, 28.97, 27.22, 27.15, 23.9, 22.6, 14.1 ppm.

*Synthesis of (±,Z)-10-Heptadecen-2-yl acetate*.

(*Z*)-10-Hexadecen-2-one (100 mg, 0.4 mmol) was dissolved in 20 ml of Ethanol and 15 mg NaBH_4_ (0.4 mmol) were added. The mixture was stirred at room temperature and after 45 minutes 5ml sat. NH_4_Cl solution was added. The mixture was extracted with diethylether (3x15 ml). The combined organic layers were washed with water (20 ml) and brine (20 ml), dried with anhydrous NaSO_4_, filtered and concentrated in vacuo to obtain crude (*Z*)-10-Heptadecen-2-ol. The alcohol was dissolved in 10 ml CH_2_Cl_2_ and 100 µl Ac_2_O, 100 µl NEt_3_ and 2 crystals of DMAP were added. The mixture was stirred at room temperature overnight, quenched with 10 ml water and extracted with diethylether (3x15 ml). The combined organic layers were washed with brine (20 ml), dried with anhydrous NaSO_4_, filtered and concentrated in vacuo. Purification via silica gel column chromatography (12:1 hexane/EtOAc) yielded racemic (*Z)-10*-Heptadecen-2-yl acetate as colourless oil (115 mg, 97%). ^1^H-NMR: (400 MHz, CDCl_3_) δ = 5.35 (m, 2H), 4.88 (hex, *J* = 6.3 Hz, 2H), 2.03 (s, 3H), 2.01 (q, *J* = 6.2 Hz, 4H), 1.51 (m, 2H), 1.29 (m, 18H), 1.20 (d, *J* = 6.4 Hz, 3H), 0.89 (t, *J* = 6.8 Hz, 3H) ppm; ^13^C-NMR: (100 MHz, CDCl_3_) δ = 170.8, 130.0, 129.8, 71.1, 35.9, 31.8, 29.7, 29.4, 29.2, 29.0, 27.22, 27.15, 25.4, 22.6, 21.4, 19.9, 14.1 ppm; EI-MS [m/z (relative intensity)]: 41 (58), 43 (100), 54 (45), 55 (84), 67 (63), 68 (51), 69 (57), 81 (67), 82 (89), 83 (40), 95 (63), 96 (77), 97 (23), 109 (33), 110 (38), 124 (49), 138 (38), 152 (12), 166 (8), 180 (5), 194 (7), 236 (35), 281 (4 [M-CH_3_]^+^).

*Separation of (R)- and (S,Z)-10-Heptadecen-2-yl acetate*.

100 mg of (±,*Z*)-10-Heptadecen-2-yl acetate were dissolved in 1ml of acetone and 7 ml of phosphatebuffer (0,1M, pH = 7.0) were added to create a fine suspension. Immobilized lipase from *Candida antarctica* was added (40 mg, 2,9 U/mg) and the mixture was stirred at room temperature. The reaction was monitored via GC-MS using a chiral column (Cyclodex B, Agilent). After 20 h the mixture was extracted with diethylether (3x10 ml), washed with brine (10 ml), dried with anhydrous NaSO_4_, filtered and concentrated in vacuo. Purification via silica gel column chromatography (9:1 hexane/EtOAc) yielded pure (*R,Z*)-10-Heptadecen-2-ol (27 mg) and (*Z*)-10-Heptadecen-2-yl acetate (67 mg of a 3:1 mixture of the (*S*)- and (*R*)-enantiomer). (*R,Z*)-10-Heptadecen-2-ol was reacetylated using the same protocol as described before to give (*R,Z*)-10-Heptadecen-2-yl acetate (25mg, ee = 99%, optical rotation: [α]^20^_589_ = +1.3, 3.01 mg/ml hexane). The remaining mixture of the (*S*) and (*R*)-enantiomer was again subjected to enzymatic hydrolysis until all of the remaining (*R*)-enantiomer was hydrolysed (ca. 48h). The mixture was worked up as described before and purified via silica gel column chromatography (9:1 hexane/EtOAc) to obtain (*S,Z*)-10-Heptadecen-2-yl acetate (31 mg, ee = 97.5%, optical rotation: [α]^20^_589_ = -1.4, 3.55 mg/ml hexane ).

**Synthesis of Heptadec-2-yl acetate (HDA)**

*Synthesis of Heptadecan-2-one.*

Palmitic acid (1.6 g, 6.25 mmol) was dissolved in 25 ml of THF and cooled to -78°C. A 1.6 M solution of MeLi in diethylether (8.6 ml, 13.75 mmol) was added dropwise and the mixture was allowed to warm to room temperature by removing the cooling bath. The reaction was monitored via TLC. Upon completion (after ca. 2.5h) the mixture was poured into 40 ml of 1N HCl at 0°C and extracted with diethylether (3x40 ml). The combined organic layers were washed with saturated NaHCO_3_ (50 ml) and brine (50 ml), dried with anhydrous NaSO_4_, filtered and concentrated in vacuo. After purification via silica gel column chromatography (12:1 hexane/EtOAc) Heptadecan-2-one was obtained as white wax (1.4 g, 90%). ^1^H-NMR: (400 MHz, CDCl_3_) δ = 2.41 (t, *J* = 7.5 Hz, 2H), 2.13 (s, 3H), 1.57 (m, 2H), 1.29 (m, 26H), 0.89 (t, *J* = 6.8 Hz, 3H) ppm; ^13^C-NMR: (100 MHz, CDCl_3_) δ = 209.4, 43.8, 31.9, 29.8, 29.7, 29.7, 29.6, 29.5, 29.4, 29.3, 29.2, 23.9, 22.7, 14.1 ppm.

*Synthesis of (±)-Heptadecan-2-ol.*

Heptadecan-2-one (1.35 g, 5.3 mmol) was dissolved in 100 ml of ethanol and NaBH_4_ (0.8 g, 21 mmol) was added at 0°C. The mixture was stirred for 2h at room temperature, quenched with 50 ml of sat. NH_4_Cl-solution diluted with 50 ml water and extracted with diethylether (2x100 ml). The combined organic layers were washed with saturated NaHCO_3_ (50 ml) and brine (50 ml), dried with anhydrous NaSO_4_, filtered and concentrated in vacuo. The residue was chromatographed on silica gel (9:1 hexane/EtOAc) to yield (±)-Heptadecan-2-ol as a white wax (1.2 g, 88%). ^1^H-NMR: (400 MHz, CDCl_3_) δ = 3.78 (h, *J* = 5.9 Hz, 1H), 1.20-1.52 (m, 28H), 1.18 (t, *J* = 5.9 Hz, 3H), 0.88 (d, *J* = 6.6 Hz, 3H)ppm; ^13^C-NMR: (100 MHz, CDCl_3_) δ = 68.2, 39.4, 31.9, 29.67, 29.64, 29.61, 29.60, 29.3, 25.8, 23.5, 22.7, 14.1 ppm.

*Synthesis of (±)-Heptadec-2-yl acetate.*

(±)-Heptadecan-2-ol (1.17 g, 4.6 mmol) was dissolved in 20 ml CH_2_Cl_2_ and 940 μl Ac_2_O, 1.4 ml NEt_3_ and 20 mg of DMAP were added. The mixture was stirred at room temperature for 3h, quenched with 20 ml ice-water and extracted with diethylether (3x30 ml). The combined organic layers were washed with water (40 ml) and brine (40 ml), dried with anhydrous NaSO_4_, filtered and concentrated in vacuo. Purification via silica gel column chromatography (12:1 hexane/EtOAc) yielded (±)-Heptadec-2-yl acetate as a white wax (1.36 g, 99%). ^1^H-NMR: (400 MHz, CDCl_3_) δ = 4.88 (h, *J* = 6.3 Hz, 1H), 2.02 (s, 3H), 1.51 (m, 2H), 1.20-1.37 (m, 26H), 1.20 (d, *J* = 6.3 Hz, 3H), 0.88 (t, *J* = 6.8 Hz, 3H)ppm; ^13^C-NMR: (100 MHz, CDCl_3_) δ = 170.5, 71.1, 35.9, 31.9, 29.68, 29.65, 29.61, 29.57, 29.53, 29.45, 29.35, 25.4, 22.7, 21.4, 19.9, 14.1 ppm; EI-MS data in agreement with data from Wiley Subscription Services, Inc. (US) via SciFinder.

**Synthesis of (19*Z*,22*Z*)- Octacosadienyl acetate (OCDA)**.

*Synthesis of 3-Eicosyn-1-ol.*

Freshly prepared THP-protected 3-Butynol^2^ (2.5 g, 16.23 mmol) was added dropwise to a suspension of NaNH_2_ (1.54 g of a 50% suspension in toluene, 19.5 mmol) in THF (15 ml) at 0°C. After 1h, DMSO (15 ml) and 1-Bromohexadecane (5.4 g, 17.7mmol) were added and the mixture was stirred for 3h at room temperature. The reaction mixture was quenched with water (20 ml) and extracted with diethylether (3 x 25 ml). The combined organic phases were washed with brine and dried over MgSO_4_. After filtration, the solvent was removed via rotavap and the residue was chromatographed on silica gel (20:1 to 15:1 hexane/EtOAc) to obtain 1-[Tetrahydropyranyl)oxy]-3-eicosyne, which was immediately dissolved in 30 ml MeOH and treated with 20 mg *p*-Toluenesulfonic acid. After 1h at room temperature, 20 ml of saturated NaHCO_3_-solution were added and the mixture was extracted with diethylether (3 x 25ml). The combined organic phases were washed with water (25 ml) and brine (15 ml) and dried over MgSO_4_. After filtration, the solvent was removed via rotavap and the residue was chromatographed on silica gel (9:1 hexane/EtOAc) to obtain 3-Eicosyn-1-ol (2.1 g, 44% yield over two steps). ^1^H-NMR: (400 MHz, CDCl_3_) δ = 3.67 (t, *J* = 6.3 Hz, 2H), 2.41 (tt, *^3^J* = 9.3 Hz, *^5^J* = 2.4 Hz, 2H), 2.14 (tt, *^3^J* = 10.6 Hz, *^5^J* = 2.4 Hz, 2H), 1.48 (m, 2H), 1.30 (m, 26H), 0.89 (t, *J* = 6.8 Hz, 3H)ppm; ^13^C-NMR: (100 MHz, CDCl_3_) δ = 82.5, 76.3, 61.3, 31.9, 29.7, 29.6, 29.5, 29.4, 29.3, 29.2, 29.1, 29.0, 28.9, 23.1, 22.6, 18.7, 14.0 ppm.

*Synthesis of 19-Eicosyn-1-ol.*

Ethylene-1,2-diamine (11 ml) was cooled to 0°C and 1.02 g NaH (60% in mineral oil, 25.5 mmol) were added. The mixture was allowed to warm to room temperature and stirred for 2 h. The violet mixture was heated to 60°C and stirred for 1h, before it was cooled to 40°C. At this temperature 3-eicosyn-1-ol (1.2 g, 4 mmol) was added. The green reaction mixture was then heated to 70°C and stirred for 5 h. After cooling to 0°C HCl_aq_ (0.5 M, 20 ml) was added very carefully. The mixture was poured into a separatory funnel and HCl_aq_ (1 M, 20ml) was added before extracting with diethylether (3 x 20 ml). The combined organic phases were washed with HCl_aq_ (1 M, 30ml) and brine (30 ml), dried over Na_2_SO_4_ and concentrated under vacuum. Purification was done via column chromatography on silica gel (9:1 *n*-hexane/EtOAc) and 19-Eicosyn-1-ol (780 mg, 65%) was obtained as a white waxy solid. ^1^H-NMR: (400 MHz, CDCl_3_) δ = 3.64 (t, *J* = 6.6 Hz, 2H), 2.17 (td, *^3^J* = 10.3 Hz, *^4^J* = 2.6 Hz, 2H), 1.94 (t, *J* = 2.7 Hz, 1H), 1.54 (m, 4H), 1.30 (m, 28H) ppm; ^13^C-NMR: (100 MHz, CDCl_3_) δ = 84.8, 68.0, 63.1, 32.8, 29.7, 29.6, 29.5, 29.4, 29.3, 29.1, 28.8, 28.5, 25.7, 18.4 ppm.

*Synthesis of 19,22-Octacosadiyn-1-ol.*

To a suspension of CuI (1.01g, 5.3 mmol), NaI (800 mg, 5.3 mmol) and K_2_CO_3_ (554 mg, 4 mmol) in DMF (10 ml), 1-bromo-2-octyn (550 mg, 2.91 mmol) and 19-Eicosyn-1-ol (780 mg, 2.65 mmol) were added. The suspension was stirred overnight and filtered over Celite®. The filtrate was poured into 20 ml of sat. NH_4_Cl and extracted with diethylether (3 x 30 ml). The combined organic phases were washed with brine (30 ml), dried over Na_2_SO_4_ and concentrated under vacuum. Purification was done via column chromatography on silica gel (12:1 *n*-hexane/EtOAc) and *19*,*22*-Octacosadiyn-1-ol (737 mg, 69%) was obtained as a white waxy solid. ^1^H-NMR: (400 MHz, CDCl_3_) δ = 3.63 (t, *J* = 6.5 Hz, 2H), 3.12 (quin, *^5^J* = 2.4 Hz, 2H), 2.14 (tt, *^3^J* = 10.5 Hz, *^5^J* = 2.3 Hz, 4H), 1.52 (m, 6H), 1.30 (m, 32H), 0.90 (t, *J* = 6.4 Hz, 3H) ppm; ^13^C-NMR: (100 MHz, CDCl_3_) δ = 80.5, 74.5, 63.0, 32.8, 31.1, 29.7, 29.6, 29.5, 29.4, 29.3, 29.1, 29.0, 28.9, 28.8, 28.5, 25.7, 22.2, 18.72, 18.70, 13.9, 9.7 ppm.

*Synthesis of 19,22-Octacosadiynyl acetate.*

19,22-Octacosadiyn-1-ol (435 mg, 1.08 mol) were dissolved in 10 ml dichloromethane. Acetic anhydride (234 μl, 2.2 mmol), triethylamine (345 μl, 2.2 mmol) and a few crystals of 4-dimethylaminopyridine were added and the solution stirred for 2 h. Water (10 ml) was added and the mixture was extracted with diethylether (3 x 15 ml). The combined organic phases were washed with brine (30 ml), dried over Na_2_SO_4_ and concentrated under vacuum. Purification was done via column chromatography on silica gel (25:1 *n*-hexane/EtOAc) and 19,22-Octacosadiynyl acetate (435 mg, 90%) was obtained as a white waxy solid. ^1^H-NMR: (400 MHz, CDCl_3_) δ = 4.05 (t, *J* = 6.8 Hz, 2H), 3.11 (quin, *^5^J* = 2.4 Hz, 2H), 2.14 (tt, *^3^J* = 10.7 Hz, *^5^J* = 2.3 Hz, 4H), 2.04 (s, 3H), 1.61 (quin, *J* = 7.0 Hz, 2H), 1.48 (m, 4H), 1.30 (m, 32H), 0.89 (t, *J* = 6.7 Hz, 3H) ppm; ^13^C-NMR: (100 MHz, CDCl_3_) δ = 171.2, 80.4, 74.5, 64.6, 31.1, 29.7, 29.6, 29.5, 29.4, 29.3, 29.1, 28.9, 28.8, 28.6, 28.5, 25.9, 22.2, 21.0, 18.72, 18.70, 13.9, 9.7 ppm.

*Synthesis of (19Z,22Z)-Octacosadienyl acetate.*

19,22-Octacosadiyn-1-ol acetate (200 mg, 0.44 mmol) was dissolved in 4 ml MeOH and hydrogenated in an H_2_-atmosphere in the presence of 15 mg Lindlar catalyst (Sigma). The reaction was monitored via GC-MS and after completion (ca. 15 h) the mixture was diluted with diethylether (15 ml), filtered over Celite® and concentrated in vacuum. Purification was done via reversed-phase column chromatography on C18-silica gel (3:1 MeOH/CHCl_3_) to obtain 19*Z*,22*Z*-Octacosadienyl acetate (180 mg, 91%) as a white waxy solid. ^1^H-NMR: (400 MHz, CDCl_3_) δ = 5.35 (m, 4H), 4.05 (t, *J* = 6.8 Hz, 2H), 2.77 (t, *J* = 6.4 Hz, 2H), 2.05 (q, *J* = 6.8 Hz, 4H), 2.04 (s, 3H), 1.61 (quin, *J* = 7.0 Hz, 2H), 1.30 (m, 32H), 0.89 (t, *J* = 6.8 Hz, 3H) ppm; ^13^C-NMR: (100 MHz, CDCl_3_) δ = 171.2, 130.2, 128.0, 64.7, 31.5, 29.7, 29.6, 29.5, 29.4, 29.3, 29.2, 28.6, 27.25, 27.20, 25.9, 25.6, 22.6, 21.0, 14.1 ppm; EI-MS [m/z (relative intensity)]: 54 (25), 55 (100), 56 (22), 57 (41), 61 (22), 67 (50), 68 (41), 69 (78), 70 (17), 71 (18), 80 (13), 81 (53), 82 (80), 83 (71), 84 (10), 85 (10), 95 (50), 96 (78), 97 (52), 109 (31), 110 (34), 111 (22), 123 (20), 124 (21), 137 (13), 138 (14), 152 (12), 390 (37), 391 (11), 448 (17 [M]^+^).

1	Francis, G. W. Alkylthiolation for the Determination of Double-Bond Position in Unsaturated Fatty-Acid Esters. *Chem Phys Lipids* **29**, 369-374, doi:Doi 10.1016/0009-3084(81)90070-0 (1981).

2	Rao, A. V. R. & Reddy, D. R. A Simple Methodology for the Synthesis of 3,4-Dihydroisocoumarins. *Synthetic Commun* **16**, 97-102, doi:Doi 10.1080/00397918608057694 (1986).
